# StartLink+: Prediction of Gene Starts in Prokaryotic Genomes by an Algorithm Integrating Independent Sources of Evidence

**DOI:** 10.1101/2020.10.25.352625

**Authors:** Karl Gemayel, Alexandre Lomsadze, Mark Borodovsky

## Abstract

Algorithms of *ab initio* gene finding were shown to make sufficiently accurate predictions in prokaryotic genomes. Nonetheless, for up to 15-25% of genes per genome the gene start predictions might differ even when made by the supposedly most accurate tools. To address this discrepancy, we have introduced StartLink+, an approach combining *ab initio* and multiple sequence alignment based methods. StartLink+ makes predictions for a majority of genes per genome (73% on average); in tests on sets of genes with experimentally verified starts the StartLink+ accuracy was shown to be 98-99%. When StartLink+ predictions made for a large set of prokaryotic genomes were compared with the database annotations we observed that on average the gene start annotations deviated from the predictions for ~5% of genes in AT-rich genomes and for 10-15% of genes in GC-rich genomes.

## Introduction

Accurate gene finding creates a solid foundation for downstream inference such as construction of the species proteome, functional annotation of proteins, prediction of cellular networks, etc. Gene starts determine boundaries for the gene upstream regions containing signals regulating gene expression (Stormo et al. 1982; de Boer and Hui 1990; Resch et al. 1996; Laursen et al. 2005).

Gene starts can be experimentally determined by several methods, such as N-terminal protein sequencing (Sazuka et al. 1999; Rudd 2000; Yamazaki et al. 2006; Aivaliotis et al. 2007; Lew et al. 2011; Zhou and Rudd 2013; de Groot et al. 2014), mass-spectroscopy (Rison et al. 2007), frame-shift mutagenesis (Smollett et al. 2009). Still the total number of genes with experimentally verified starts is limited i.e. 2,443 such genes (Hyatt et al. 2010) and 2,925 genes (Lomsadze et al. 2018) (from 7-10 different species were used in benchmarking of gene finding algorithms.

Current state-of-the-art tools largely agree with each other when tested on a (limited) set of genes with experimentally verified gene-starts and achieve high accuracy. However, they show striking disagreements on larger sets of genes. Here, we describe a new approach for predicting gene-starts that combines *ab initio* and multiple sequence alignment based methods; this approach makes gene start predictions for 73% of genes (on average) per genome with average error rate observed to be less than 1%, as it was assessed on the sets of genes with verified gene starts. Comparisons with existing *ab initio* tools as well as with database annotations suggest that the new approach could provide justification for revision of current gene start annotation in up to 15% of genes in a genome.

## Motivation

It was observed that when Prodigal and GeneMarkS-2 were run on sets of genes with experimentally verified genes (2,925 genes in total), they predicted gene starts with average accuracy of ~95% (Lomsadze et al. 2018). We also observed that these two tools disagree in gene start predictions, in the same verified sets, in ~6% of genes on average.

In a computational experiment with a set of 5,488 representative prokaryotic genomes (Fig. 1) we have compared gene start predictions made by GeneMarkS-2 (Lomsadze et al. 2018), by Prodigal (Hyatt et al. 2010), and by the PGAP pipeline (Tatusova et al. 2016) guided by alignments of annotated starts of homologous genes. We observed that in many genomes, especially those with high GC content, gene start predictions may differ from annotations on average for 7-22% of the genes in a given genome.

**Figure 1:**
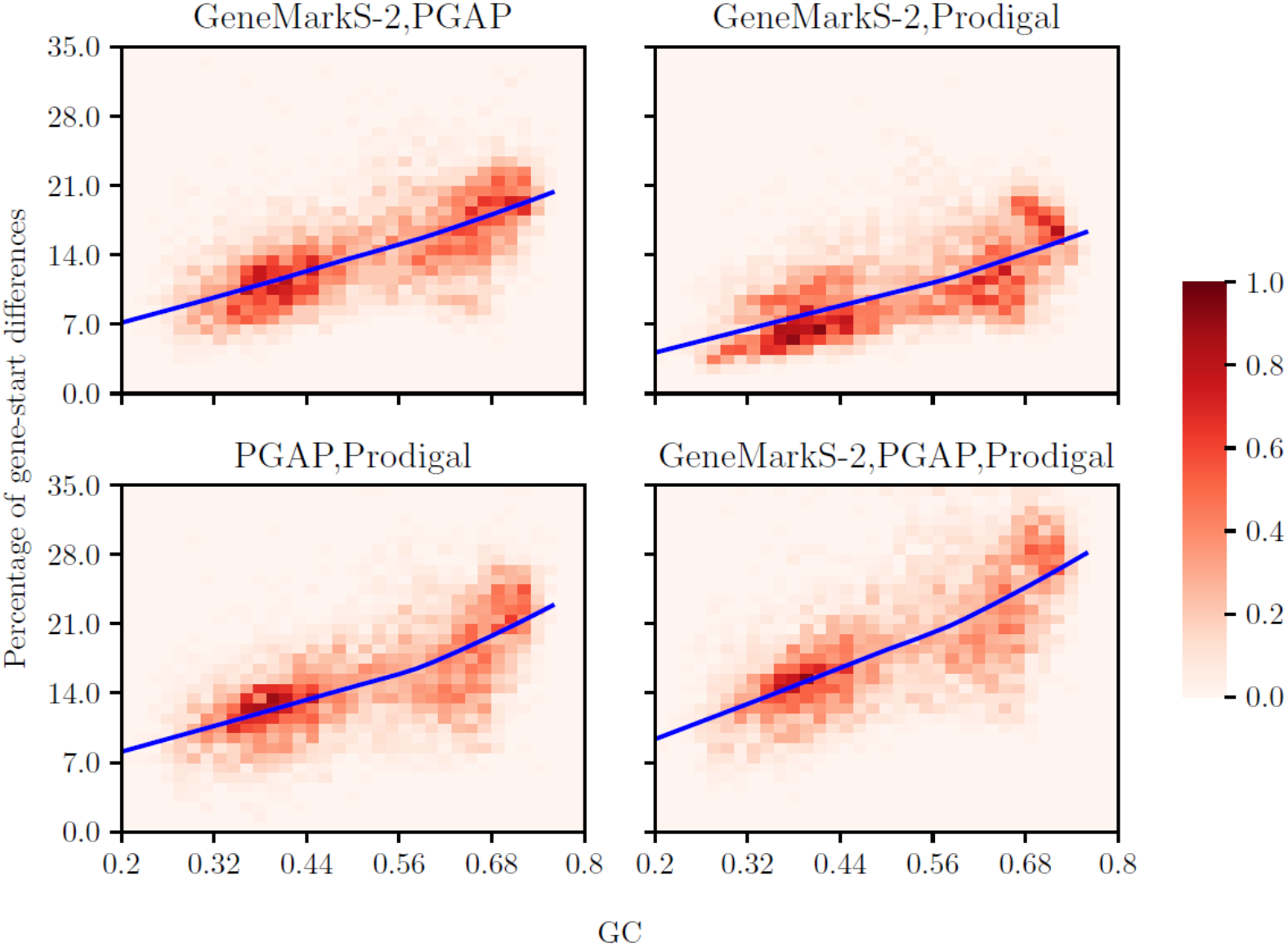
Disagreements of the three tools (Prodigal, GeneMarkS-2, and NCBI’s PGAP) in gene start predictions observed in experiments with the NCBI collection of 5,488 representative genomes. The percentage of differences in gene start predictions (per genome) was computed by taking the number of genes where at least one of the tools has a difference in gene starts to the other(s), divided by the number of genes in a given genome having predictions by all the tools (related to a particular graph). Each colored square shows the percentage of genomes within a GC bin (from the total number of 5,488) that have a certain percentage of gene-start differences. The color of the square indicates the percentage value; the color coding key is given by the bar on the right, ranging from 0 to 1%. The average percentage of genes (per genome) with difference in gene start predictions by particular tools is shown by a solid line as a function of GC content.

The differences in gene start predictions could appear due to complexity and variability of sequence patterns in the genes’ upstream regions. GeneMarkS (Besemer et al. 2001) and Prodigal were able to infer ribosome binding sites (RBS) models with patterns different from the canonic Shine-Dalgarno (SD) RBS (Shine and Dalgarno 1974; Barrick et al. 1994). Frequent in archaea leaderless mRNAs, first discovered in *Pyrobaculum aerophilum* (Slupska et al. 2001), don’t leave space RBS upstream of start codons of many first genes in operons. Knowledge of genes with leaderless transcription is instrumental in pathogen studies as some antibiotics inhibit translation initiation on leadered transcripts and not on leaderless ones (Brandi et al. 2006; Schuwirth et al. 2006; Kaberdina et al. 2009; Muller et al. 2016; Lange et al. 2017; Sawyer et al. 2018). In absence of RBS sites, the promoter signal (located then on a close distance to the translation start site) could be used to improve gene-start detection. Still, Prodigal is primarily oriented to search for an SD RBS, and its core RBS model was optimized on the *Escherichia coli* genes with verified starts (Rudd 2000; Hyatt et al. 2010). Both GeneMarkS and Prodigal have difficulty to work correctly in genomes where leaderless transcription and RBS-based translation are present at the same time.

More recently developed GeneMarkS-2 created models for multiple gene expression mechanisms within the same genome. We have found from analysis of 5,007 representative genomes (238 archaea and 4,769 bacteria) that 16.4% of archaeal and 61.5% of bacterial genomes predominantly used the SD-RBS. The remaining 83.6% of archaea were predicted to frequently use leaderless transcription (along with SD-RBS for some genes). These predictions for archaeal genomes were supported by experimental findings, e.g. for *Halobacterium salinarum*, *Haloferax volcanii*, and *Thermococcus onnurineus* (Koide et al. 2009; Babski et al. 2016; Cho et al. 2017). On the other hand, from the remaining 38.5% of bacterial species 10.4% were found to use a non-SD type RBS (e.g. *Bacteroides* (Wegmann et al. 2013)), 21.6% use leaderless transcription (often for up to 40% of transcripts in a genome, e.g. *Mycobacterium tuberculosis* (Cortes et al. 2013; Shell et al. 2015; Gualerzi and Pon 2015; Nakagawa et al. 2017)), and the remaining 6.5% use an SD-RBS for a small fraction of genes while the majority had a signal with a very weak sequence pattern, that indicates an unknown mechanism of translation initiation (e.g. *Cyanobacteria* (Mutsuda and Sugiura 2006)).

In this work, we developed a multiple sequence alignment based gene start prediction algorithm called *StartLink*. We have not used existing gene start annotations as well as information on sequence patterns related to translation-initiation mechanism, such as RBS or promoter site models. As an evidence, however, we have used results of alignment of multiple unannotated syntenic genomic sequences containing genes extended to the longest open-reading-frames (LORFs). By design, StartLink could serve as a standalone predictor of gene starts. It is applicable for all the genes that have sufficient number of homologs. Particularly, it is applicable for finding starts of genes residing in short contigs (e.g. assembled from metagenomics reads) for which GeneMarkS-2 (and other whole genome *ab initio* gene finders) may not perform well due to insufficient volume of sequence data that could be used for supervised or unsupervised training.

We have demonstrated that if StartLink and GeneMarkS-2 predicted the same gene start, such a prediction had an error rate of 1-2%. When GeneMarkS-2 and StartLink predictions coincide we designate them as StartLink+ predictions.

## Datasets

As of November 4, 2019, NCBI’s RefSeq database had over 183,689 annotated prokaryotic genomes. To reduce time for search for homologs, the search space could be limited to a clade the query species belongs to (e.g. see Table 1). Among genomes with the same taxonomy ID we selected the one with the most recent annotation date. All longest open reading frames (LORFs) of annotated genes in the selected genomes were extracted, translated and a BLASTp database was built.

**Table 1.**
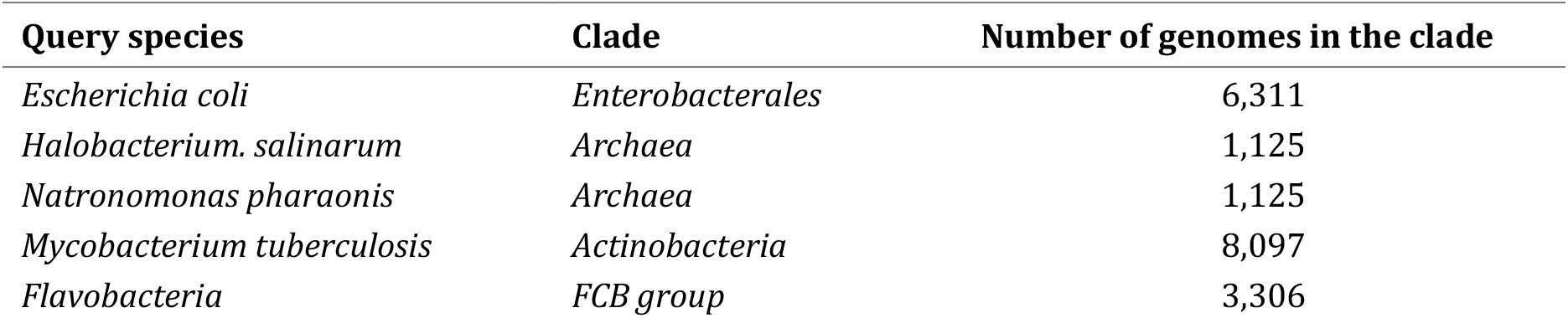
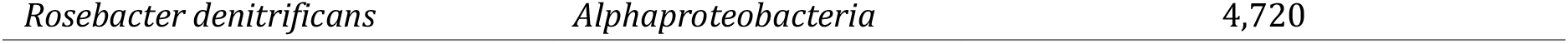
Reference clades for the five query species

| Query species | Clade | Number of genomes in the clade |
| --- | --- | --- |
| *Escherichia coli* | *Enterobacterales* | 6,311 |
| *Halobacterium. salinarum* | *Archaea* | 1,125 |
| *Natronomonas pharaonis* | *Archaea* | 1,125 |
| *Mycobacterium tuberculosis* | *Actinobacteria* | 8,097 |
| *Flavobacterium* | *FCB group* | 3,306 |

The five species, bacteria *E. coli* (Rudd 2000; Zhou and Rudd 2013), *M. tuberculosis* (Lew et al. 2011), and *R. denitrificans* (Bland et al. 2014), and archaea *H. salinarum* and *N. pharaonis* (Aivaliotis et al. 2007) listed in Table 1 had, as of December 2019, the largest numbers of genes with starts verified by N-terminal sequencing (see Table 3):

We have conducted experiments with genomes from four clades (with the numbers of randomly selected genomes in parenthesis): *Archaea* (97), *Actinobacteria* (95), *Enterobacterales* (106), and *FCB group* (96). The clades selection was guided by the study of patterns in upstream regulatory regions (Lomsadze et al. 2018). Archaeal genomes have large numbers of genes with leaderless transcription. Clade *Actinobacteria* has predominantly high-GC genomes with a significant number of genes with leaderless transcription. The *Enterobacterales* clade has mostly mid-GC genomes that carry genes with an RBS of the Shine-Dalgarno type. Finally, the *FCB group* has low-to-mid-GC genomes that carry genes with a ‘non-canonical’ AT-rich RBS (Lomsadze et al. 2018).

## Methods

### Metrics for gene-start prediction performance

Given a reference set of genes, set *G*, we consider its subset S for which a particular algorithm predicts gene starts. We define *accuracy*, Acc(S,G), *error rate*, Err(S,G), and *coverage*, Covr(S,G), by the following formulae:

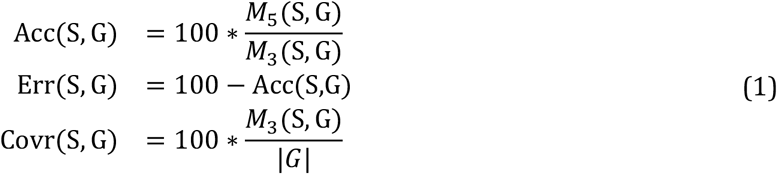

Here *M*_5_(S, G) and *M*_3_(S, G) are the numbers of genes in *S* that match genes in *G* by both 5’ and 3’ ends, and by only 3’ ends respectively.

### StartLink

Our task is to identify the start codon of a prokaryotic gene within its longest open reading frame (LORF) embedded in sequence Q (query). The StartLink algorithm uses cross-species syntenic genomic sequences upon making the following three steps (Fig. 2).

1. Gathering a set of target genomic sequences with Q as a query in the search.
2. Eliminating evolutionarily too close and too remote to Q target sequences as well as target sequences too close to each other and constructing a multiple sequence alignment (MSA).
3. Selecting gene start among possible candidates within the LORF in Q.

**Fig. 2:**
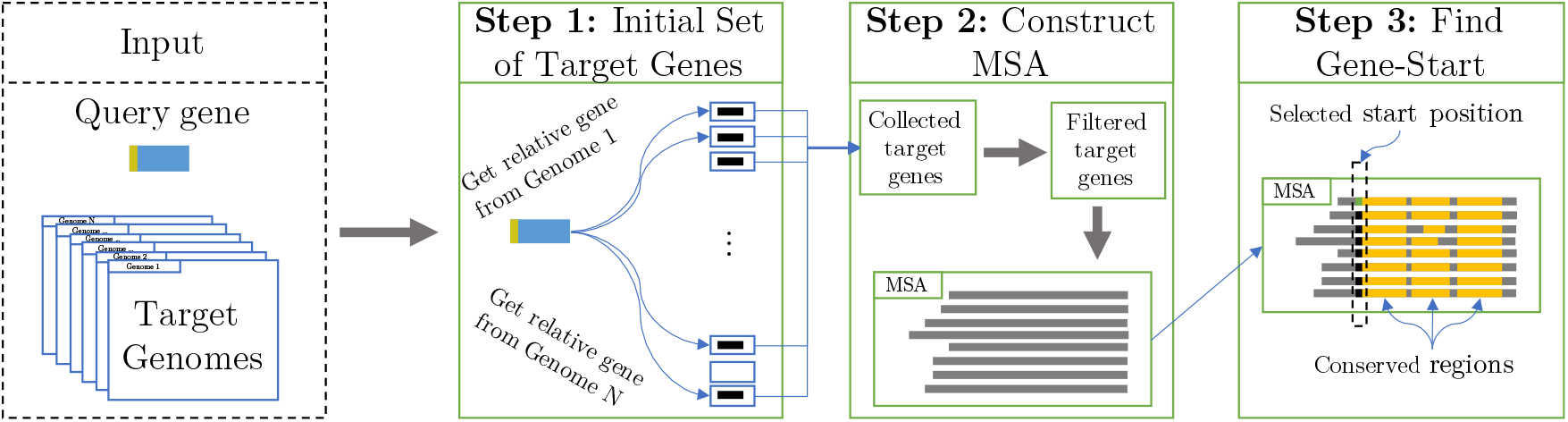
A high-level schematic of StartLink (Fig. 3 shows details of Step 3).

#### Step 1: Finding homologs

A protein product of a query gene is used in Diamond BLASTp (Buchfink et al. 2015) to find in a database of target proteins (described in more detail further down) a set of target proteins (and genes) that have significant similarity to the query. We remove any target whose protein alignment with the query does not cover 80% of either the query or target sequences. This step helps eliminate, among others, the longer target proteins where a query protein does not align close to the target gene-start.

#### Step 2: Filtering of targets and constructing the MSA

Having the set of target proteins and their genes we proceed to build an informative MSA for gene-start inference. Each target gene is extended to LORF and translated into amino acid sequence. The protein MSA is constructed by the Clustal Omega algorithm (Sievers and Higgins 2018) from 50 randomly selected translated LORFs along with the translated LORF of the query. Next, the algorithm constructs protein alignment guided alignments of nucleotide sequences and computes the query-to-target and target-to-target evolutionary distances according to the Kimura 2-parameter model (Kimura 1980):

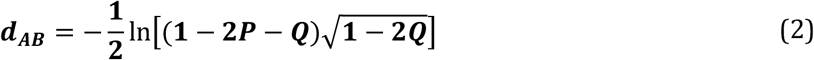

Here *P* and *Q* are the fractions of positions in the pairwise alignment of DNA sequences which pairs of nucleotides exhibit transition or transversion mutations, respectively. We retain target sequences that fall inside the *d*_*AB*_ range [0.1, 0.5] with respect to the query. This distance range was selected based on the tests and the analysis of MSAs obtained from randomly selected genomes (see Supplementary Material).

The Kimura distance is usually computed for a global alignment of two DNA sequences. We have observed that for closely related genomic sequences, a local alignment (derived from the readily available BLASTp output) could provide sufficiently accurate distance value, thus saving the effort of realigning sequence pairs (see Supplementary Note 6).

Next, we reduce the set of sequences selected in the *d*_*AB*_ range [0.1,0.5] due to the following considerations:

1. If two syntenic sequences are too similar, then one of them is redundant and could be removed.
2. We expect that aligned syntenic genomic sequences would carry conserved patterns downstream from true gene starts.
3. Elimination of rather distant sequences would significantly improve the alignment, as their presence would disrupt the pattern of conservation by insertion of a large number of gaps in MSA downstream from the gene start.

Each time a sequence is removed, we regenerate the MSA using the remaining sequences and re-apply the filtering steps. The number of sequences in the final MSA varies from 10 to 50 (see Fig. 16); the average number of MSA sequences is clade specific. We observed that MSAs with low numbers of targets (e.g. about 10) still contain informative sequences.

#### Step 3: Identification of the gene start in the query sequence

The algorithm predicts the gene start by analyzing patterns of conservation in the MSA in one of the three following steps.

##### Step A: Search for conserved blocks in protein MSA and the simplest case of the gene start identification

Given a protein MSA constructed from the translated LORFs of a query and its targets, the algorithm searches for the left-most *block with high conservation score* (see below). Notably, we assume that the nucleotide sequences of the true genes in the corresponding set of nucleotide sequences do not overlap with the upstream genes. If a left-most protein block with high score is detected and there is only one gene start candidate in the nucleotide query upstream of the block, this candidate is predicted to be the gene-start. Otherwise, the algorithm proceeds to step B. Note that the start assignment in step A does not require conservation of the start candidate itself.

For a protein MSA block of length *r* (where *r* = 10*aa* not including possible N-terminal), a conservation (identity) score is computed by the formula

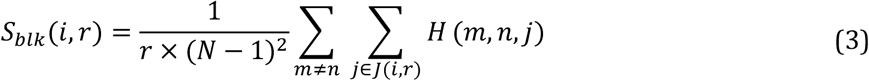

where *J*(*i*) is the set of *r* positions downstream of position *i*, with no gap in the query; *H*(*m*, *n*, *j*) is 1 if and only if sequences *m* and *n* match each other at position *j* in the alignment, and *N* is the total number of sequences in MSA. A block with *S*_*blk*_(*i*, *r*) larger than 0.5 is identified as conserved. This threshold corresponds to uninformed, majority-vote approach, a reasonable option when little ground-truth data is available.

##### Step B: Identification of the gene start in the presence of overlapping genes

If a query LORF overlaps with the 3’ end of the upstream gene (which is relatively easy to reliably determine), such an overlap is likely to appear in syntenic sequences (at a sufficiently close evolutionary distance, see Figs. 8 and 9). It was observed that ATG, GTG, or TTG codons of a LORF situated near the 3’ end of the upstream gene have elevated frequency of being true starts (Lukashin and Borodovsky 1998; Huber et al. 2019). It is plausible that the ribosome can efficiently reassemble at such a gene start upon completing the translation of the upstream gene. Therefore, StartLink attempts to identify a conserved in the MSA gene-start candidate within 9 *nt* distance near the 3’ end of the upstream gene. The conservation score for the candidate with MSA position *i* is defined by the fraction of targets that have a gene start candidates within 6*nt* distance from position *i*. Formally, the identity score for position *i* is defined as

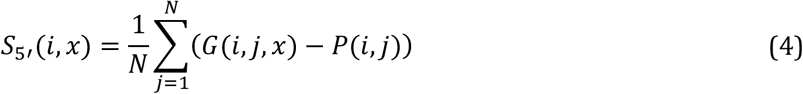

where

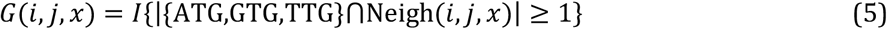

Here, *I*{⋅} is the indicator function, |⋅| computes the size of a set, and Neigh(*i*, *j*, *x*) is the set of codons within a distance of *x* codons around position *i* in sequence *j*. Thus, *G*(*i*, *j*, *x*) is 1 if an ATG, GTG, or TTG exists in the neighborhood, and 0 otherwise. The term *P*(*i*, *j*) penalizes for the appearance of the codons being synonymous to GTG, or TTG, but not serving as start codons; *P*(*i*, *j*) = 1 if such a codon exists in position *i* of sequence *j* in the MSA, and 0 otherwise. *If S*_5’_(*i*, *x*) > 0.5 the candidate is selected as a predicted start otherwise the algorithm moves to Step C.

For additional justification of the use of step C we introduced the following consideration based on analysis of a large set of query genes and their homologs (targets). We found that if a query gene was overlapped by the upstream gene (in the same strand) or if the upstream intergenic region was very short (less than 10nt), then such a configuration was preserved for genes in genomes of closely related species as well. As a quantitative measure of this evolutionary conservation, we introduced the following function.

Let consider a query gene along with its targets defined by similarity search and included in the MSA (N sequences, N > 10); we have called this set *a component*. Let *D*(*n*) be the length of intergenic region from the end of the upstream gene to the start of the downstream gene, *n*, and let *x* be the most frequent *D*(*n*) observed in the component (i.e. the mode). Then, the measure of conservation of intergenic region being *x* nucleotides long is defined as

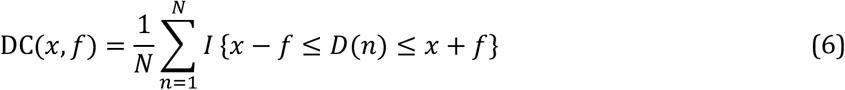

where *I*(⋅) is the indicator function and the margin *f* determines the stringency of conservation. DC value could be interpreted as a probability that in any sequence within a component the upstream gene is located *x* ± *f nt* away, where *x* is the most frequent distance in the component. The distribution of the measure of the conservation was computed and presented in the Results section (Fig. 8).

##### Step C: Identification of the gene start under ‘common’ conditions

There are two sub-steps.

###### Sub-step C-1

If in the query LORF there are multiple gene start candidates upstream to the left-most MSA block of conserved amino acids, the *S*_5’_ scores of the candidates are screened from the LORF 5’ end downstream. If a candidate has *S*_5’_ > 0.5, the algorithm moves to Step C-2. Otherwise, it moves to the next candidate. If all candidates have been exhausted, the algorithm quits without selecting any candidate as a predicted gene start.

###### Sub-step C-2

To avoid missing a true start downstream of the candidate selected in C-1 the algorithm searches for a candidate with a highest *S*_5′_ score - *S*′_5′_ in the 30nt region downstream. If *S*′_5′_ > 0.5 and if there is a conserved block (of any length up to 10 *aa*) between the two candidates, the upstream candidate is selected; otherwise, the downstream candidate is identified as the start (see Fig. 3).

**Figure 3:**
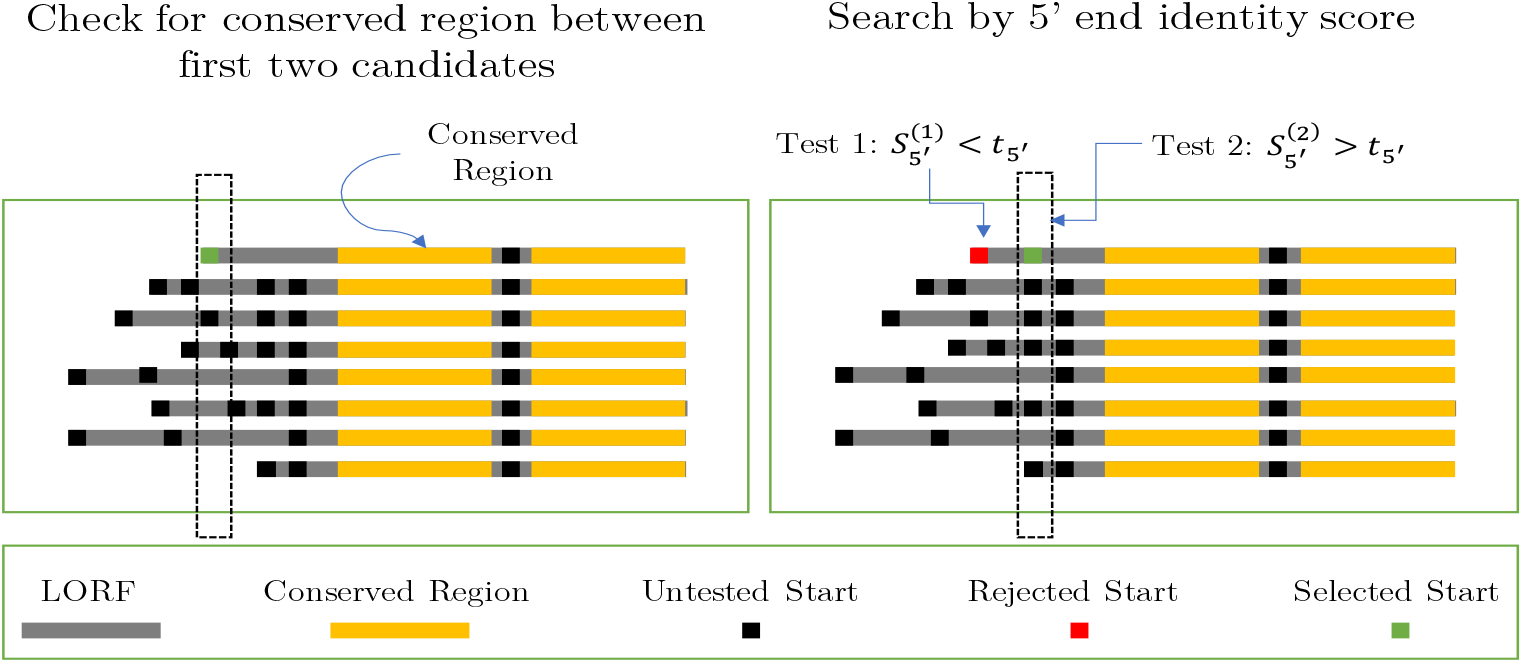
The use of MSA to identify a start of a gene in the query sequence (top sequence in each MSA). **Left panel:** Step A: the left-most conserved block is detected, with a single gene start candidate located upstream. **Right panel:** Step C: Candidate start codons are screened to find those with conservation score *S*_5′_(*i*, *x*) above a threshold *t*_5′_ = 0.5 (Supplementary Material 3)

### StartLink+: Joint use of StartLink and GeneMarkS-2

StartLink+ runs both GeneMarkS-2 and StartLink and selects genes where gene start predictions of both tools coincide. Jointly predicted starts reported as output of StartLink+. The probability of error of StartLink+ is the probability of an event that two sufficiently accurate independent tools would make exactly the same erroneous prediction. Thus, the error rate is proportional to a product of probabilities of an error of each tool. This approach may not produce predictions for all the genes in a genome, however, as we show in the results section, this approach generates gene start predictions for a significant number of genes in each genome with a chance of error close to 1% (see Table 3 and Supplementary Methods).

## Results

### StartLink+ accuracy assessment on genes with experimentally verified starts

In the genomes containing genes with verified starts we selected the genes with StartLink+ predictions. The coverage values, percentage of genes in a given set for which a particular method generates gene start predictions, are shown in Table 2 for StartLink, GeneMarkS-2 and StartLink+.

**Table 2:**
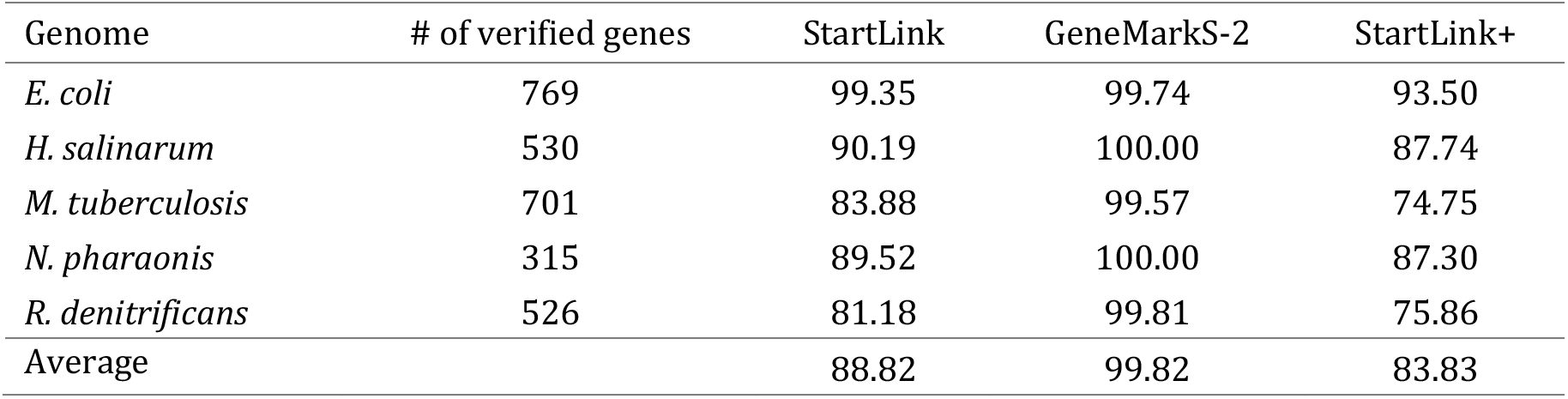
Coverage of the sets of genes with experimentally verified starts (total of 2, 841genes) by the three gene start prediction methods.

| Genome | # of verified genes | StartLink | GeneMarkS-2 | StartLink+ |
| --- | --- | --- | --- | --- |
| *E. coli* | 769 | 99.35 | 99.74 | 93.50 |
| *H. salinarum* | 530 | 90.19 | 100.00 | 87.74 |
| *M. tuberculosis* | 701 | 83.88 | 99.57 | 74.75 |
| *N. pharaonis* | 315 | 89.52 | 100.00 | 87.30 |
| *R. denitrificans* | 526 | 81.18 | 99.81 | 75.86 |
| Average |  | 88.82 | 99.82 | 83.83 |

The GeneMarkS-2 coverage deviated from 100% when the gene finder did not predict a whole gene in a particular set. StartLink was missing genes where neither A or B or C steps produced start predictions. In addition to genes missed by either GeneMarkS-2 or StartLink, StartLink+ missed genes where gene starts predicted by GeneMarkS-2 and StartLink do not match. The StartLink+ coverages observed in *M. tuberculosis* and *R. dentrificans* sets was the lowest ~ 75%.

The error rates of gene start prediction by StartLink, GeneMarkS-2 and StartLink+ were computed by comparison of the predictions with the co-ordinates of verified gene starts (Table 3). The error rates of StartLink+ were quite low. Particularly striking was the example of *M. tuberculosis*. While StartLink and GeneMarkS-2 had ~6.9% and ~9.6% error rates, respectively, the error rate of StartLink+ was ~1.3%.

**Table 3:** The error rates of StartLink, GeneMarkS-2 and StartLink+ predictions observed on the sets of genes with experimentally verified starts

| Genome | # of verified genes | Error rates |  |  |
| --- | --- | --- | --- | --- |
|  |  | StartLink | GeneMarkS-2 | StartLink+ |
| *E. coli* | 769 | 4.45 | 3.00 | 0.83 |
| *H. salinarum* | 530 | 2.73 | 1.32 | 0.43 |
| *M. tuberculosis* | 701 | 6.86 | 9.60 | 1.32 |
| *N. pharaonis* | 315 | 2.11 | 0.95 | 0.00 |
| *R. denitrificans* | 526 | 4.81 | 3.43 | 0.45 |
| Average |  | 4.19 | 3.66 | 0.61 |

Three genomes in Table 3 had high GC content: *H. salinarum* (65%), *N. pharaonis* (63%), and *M. tuberculosis* (66%). StartLink+ demonstrated a low error rate for each of these genomes (0.6%, 0.0%, 1.32%, respectively).

### Differences between predictions of StartLink+, Prodigal and PGAP

The assessment of accuracy of StartLink+ (Table 3) showed remarkably low error rates in gene-start predictions (albeit with reduction in coverage). Therefore, by application of StartLink+ to a given genome one could generate a large set of genes with vast majority of starts reliably determined. We did run StartLink+ on 443 prokaryotic genomes from four prokaryotic clades (Table 4).

**Table 4:** Number of genomes selected in each prokaryotic clade, and the number of genes with gene starts predicted by StartLink+.

| Clade | Query Genomes | StartLink | StartLink+ |
| --- | --- | --- | --- |
| *Actinobacteria* | 111 | 276,468 | 228,837 |
| *Archaea* | 108 | 243,424 | 212,162 |
| *Enterobacterales* | 119 | 451,255 | 394,271 |
| *FCB group* | 105 | 253,257 | 229,212 |
| **Total** | **443** | **1,224,404** | **1,064,482** |

Next, we compared the predicted gene starts to the PGAP annotation We observed non-uniform distribution of the percentages of genes with differences in gene start positions between PGAP and StartLink+ among the clades (Fig. 4). Particularly in *Actinobacteria* genomes, that difference reached up to 15% of genes per genome, with an average of around 10%. On the other hand, the average difference dropped to about 4.5% in *FCB group* genomes, and to ~3% in *Enterobacterales* genomes. Notably, there were inter-clade differences in average genome GC contents, e.g. *Actinobacteria* (high GC) and *Enterobacterales* (mid GC), as well as in clade-specific abundance of leaderless transcription.

**Figure 4:**
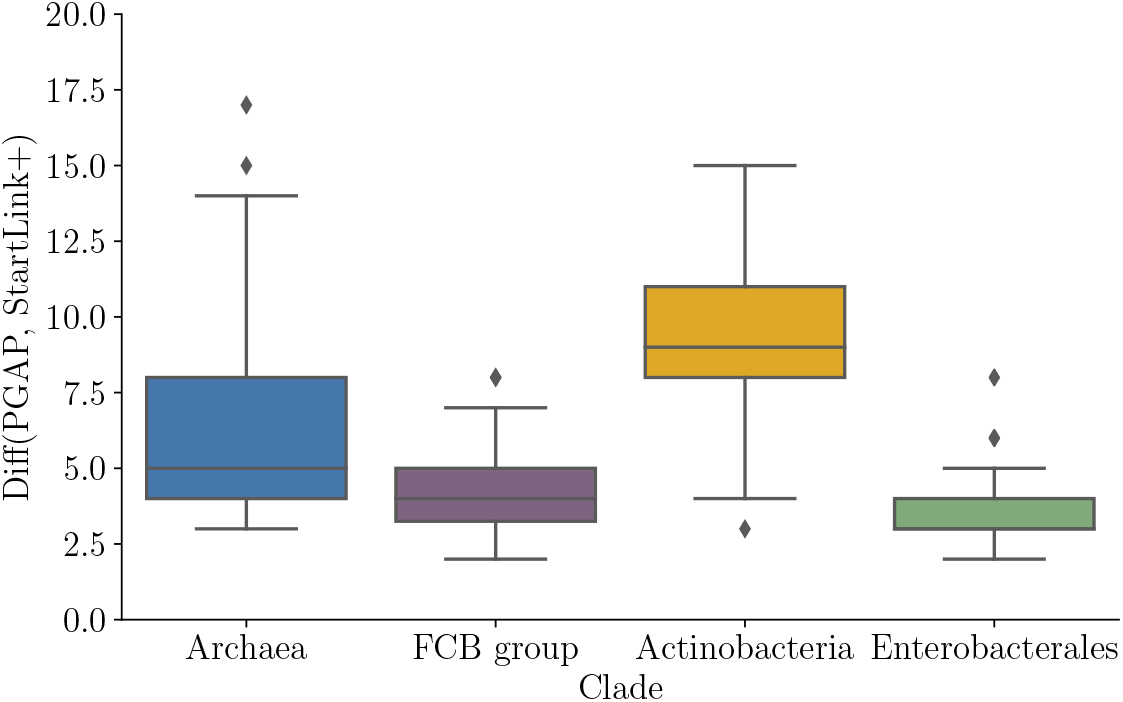
Percentages of genes with differences in gene start positions determined by PGAP and StartLink+. Distributions of average values per genome for the species of the prokaryotic clades (Table 4).

Fig. 5 shows the percentage of genes with differences in start prediction determined by PGAP and StartLink+ as well as Prodigal and StartLink+ as a function of the genome GC content. In both cases, the average percentage of gene-start differences increased as a function of GC. In *Archaea*, the percentage of differences increased in mid-GC genomes and decreased in high-GC and low-GC genomes. In *Actinobacteria*, the percentage increased across GC for both Prodigal and PGAP, but beyond 67% GC it began decreasing for Prodigal.

**Figure 5:**
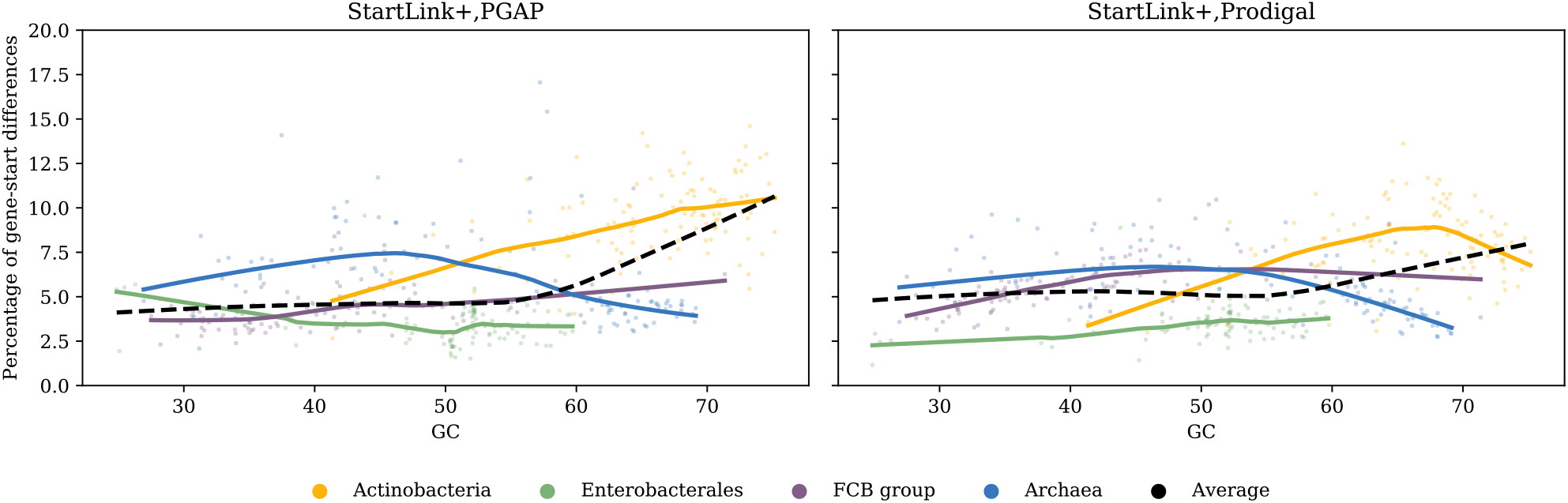
Percentage of genes per genome where predicted start positions differ between PGAP and StartLink+ (left) and Prodigal and StartLink+ (right), as a function of the genome GC content. The analysis was done for 443 genomes from four clades (Table 4)

By its design, PGAP determined many gene starts based on a conservation pattern inferred from annotated gene starts of known genes (Tatusova et al. 2016). Still, this method may not be perfect as the gene start annotation in previously sequenced genomes could be biased, also, the reference genomes may appear at different evolutionary distances. Moreover, the results of selection of reference genes in StartLink were gene specific as different genes evolve with different speed. Thus, the variations in the sets of reference genes in terms of distribution of Kimura distances might result in difference of performance in different clades.

### Gene start prediction performance per a StartLink step

As previously mentioned, StartLink output was generated at the first or second or third step (A, B or C respectively) depending on the configuration of the sequence alignment patterns. In this section, we use *sets of genes with verified starts* to assess the performance of both StartLink and StartLink+ at each step. We also show the percentage of gene-start differences (Diff) between StartLink+ prediction and PGAP annotation at each step

At step A the observed error rate was consistently low, close to 0.0% (Fig. 6, left panel). Indeed, the step A predictions were made with a strong evidence for a particular gene start; ambiguous cases were delegated to the following steps. Still, the error rates observed at steps B and C were rather low as well.

**Figure 6:**
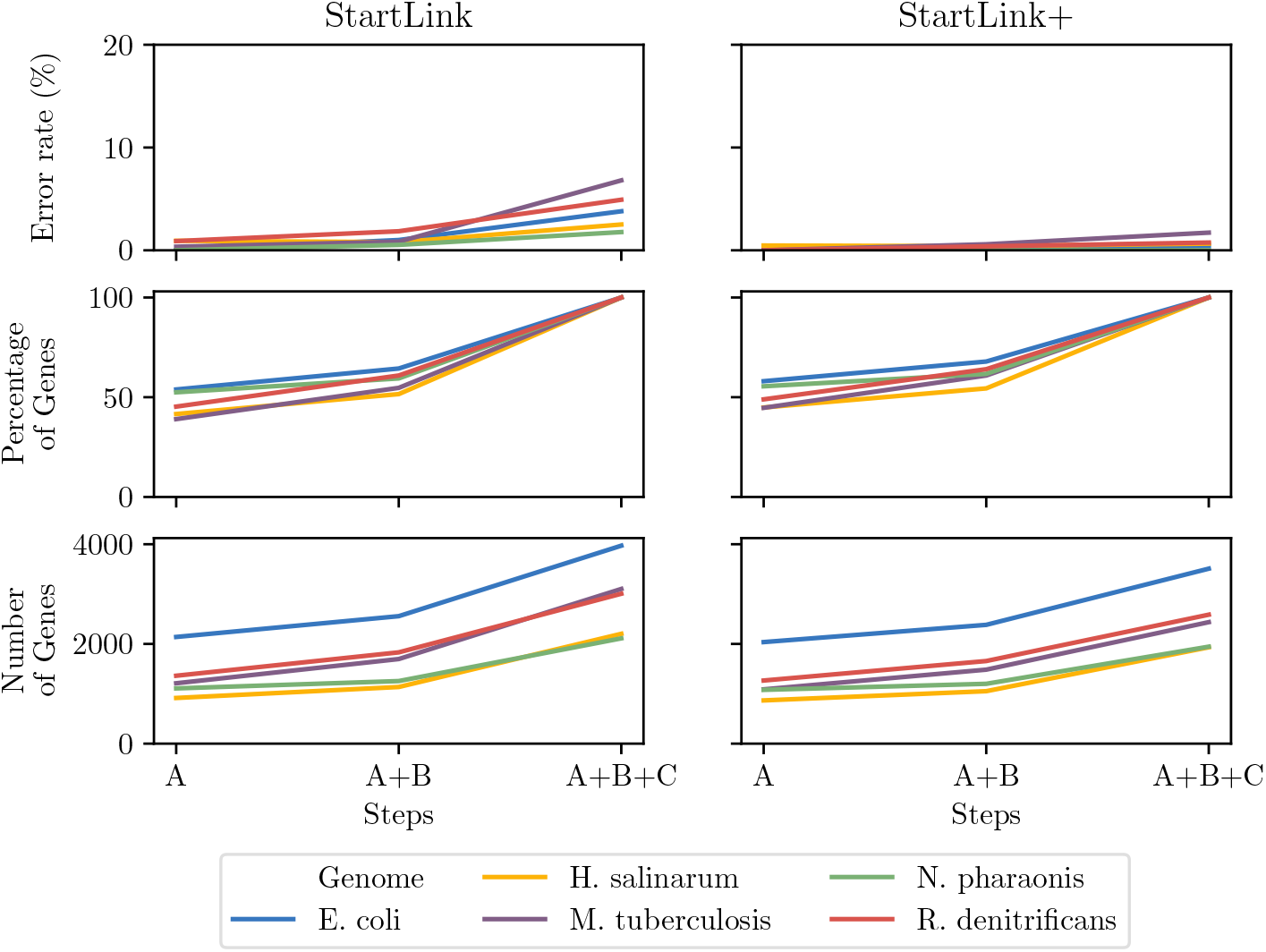
Left panels: The gene-start prediction error rate of StartLink observed at steps A, B, C on the sets of verified starts of the five species (top), the percentage of the genes in a given genome predicted by StartLink by step A alone, by steps A and B, and by all steps together (middle) and the absolute numbers of the predicted genes depicted in the middle sub-panel (bottom). Right: Same data types as in the left panel for StartLink+.

We should mention that among the genes with verified starts a very few genes had a closely situated or overlapping upstream genes. Particularly for *N. pharaonis*, the Step B was a final step for only seven genes. We also saw that StartLink+ demonstrated low error rates (Fig. 6, right panel). Interestingly, the observed pattern of differences between StartLink+ predictions and the PGAP annotation were similar but not the same in the four prokaryotic clades (Fig. 7). The differences were consistently smaller at step A (in the range of 2-6%) in comparison with steps B and C (5-12%). Similar patterns were found when comparing StartLink+ to Prodigal (data not shown).

**Figure 7:**
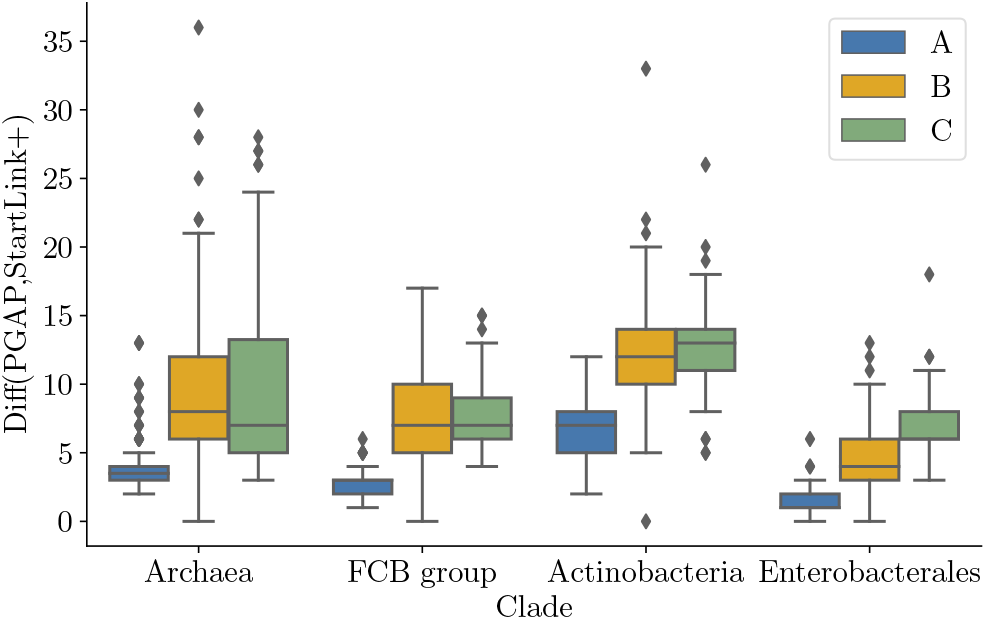
The differences between PGAP annotation and StartLink+ predictions. Comparisons were made separately for predictions made at each step of StartLink (A, B, C).

### Conservation of gene overlaps in syntenic regions

We have shown presence of conservation of overlaps and short intergenic regions defined by formula (9) for the DC measure of conservation. We have observed that the DC value increased when *x decreased* (Fig.8)

**Figure 8:**
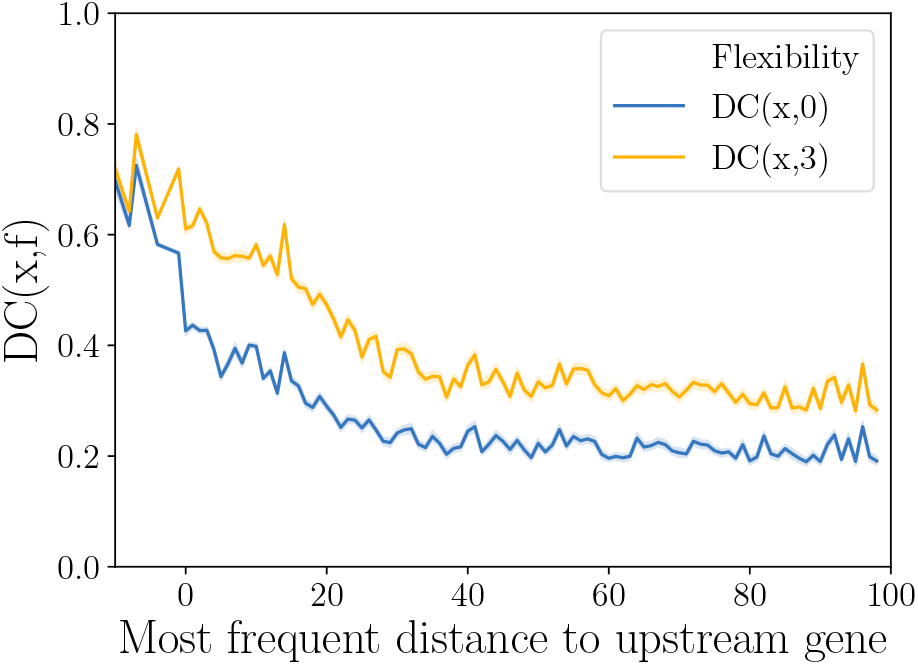
The intergenic distance conservation for f=0 and f=3 as a function of the most frequent upstream distance per component. The data was computed in experiments with 443 query genomes. The PGAP annotation was used for the positions of gene starts. The origin in X axis is situated at −9.

We saw that gene overlaps tend to be conserved within components.

To determine the percentage of components (per clade) that fell for each value of *x* we zoomed in into the range of *x nt* between −10 and 10 (Fig. 9). Most components within that range had 4 *nt* gene overlap, followed by 1 *nt* overlap. This tendency was particularly pronounced in *Actinobacteria*, where more than 60% of components had 4 *nt* gene overlap. In *FCB group*, components with 4 *nt* gene overlap constituted only 20% of all components. This decrease with respect to *Actinobacteria* could be related to the presence of AT-rich non-canonical RBS in the gene upstream regions of genomes of *FCB group* (Lomsadze et al. 2018). Such AT-rich RBS sites could have been evolved and maintained in lower GC non-coding regions rather than inside the upstream protein-coding gene with higher GC. The observed preferences for both −4 and −1 overlaps were in agreement with the previous works suggesting that gene-starts positions close to the 3’ ends of the upstream genes were favored in evolution (Lukashin and Borodovsky 1998; Huber et al. 2019).

**Figure 9:**
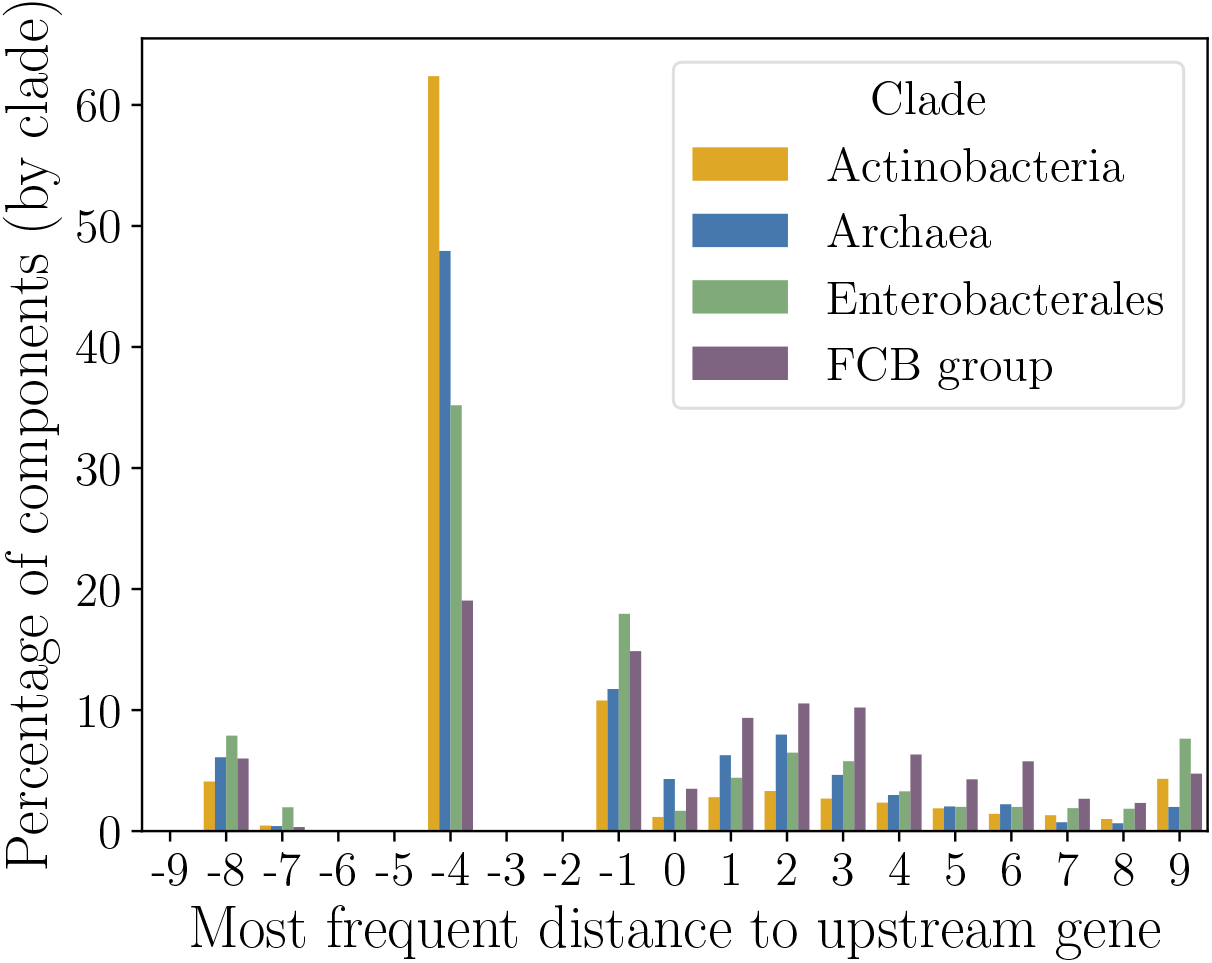
The frequency histogram of the most frequent intergenic distance *x* between same-strand genes in the MSA defined components. The *x* value is in the range from −10 to +10 nt.

### Analysis of distributions of the Kimura distances

The clade-specific accuracy of StartLink could depend of clade-specific organization of groups of homologous genes or proteins. We analyzed distributions of the Kimura distances between query genes and their targets across different clades (in the distance range [0.1, 0.5]).

Regardless of the nature of the differences in the Kimura distance distributions (caused by the variability of the speed of evolution or inhomogeneity of the database sampling), the similarity-based method, such as StartLink, had to be designed to work in a non-uniform space of homologs (Supplementary Note 2).

A set of orthologs found by similarity search for a given query *q* was filtered prior to MSA construction. This set had minimum and maximum values of the Kimura distances to the query. These two values made a vector (min K(*q*), max K(*q*)) and the frequency distribution of these vectors within the triangular space were depicted by the contour plots (Fig. 10). The plots show clear differences in the distributions of the ‘min-max’ vectors among queries in each clade. For example, most query genes in *Enterobacterales* had the ‘min-max’ vectors of the Kimura distances close to the extreme one (0.1, 0.5).

**Figure 10:**
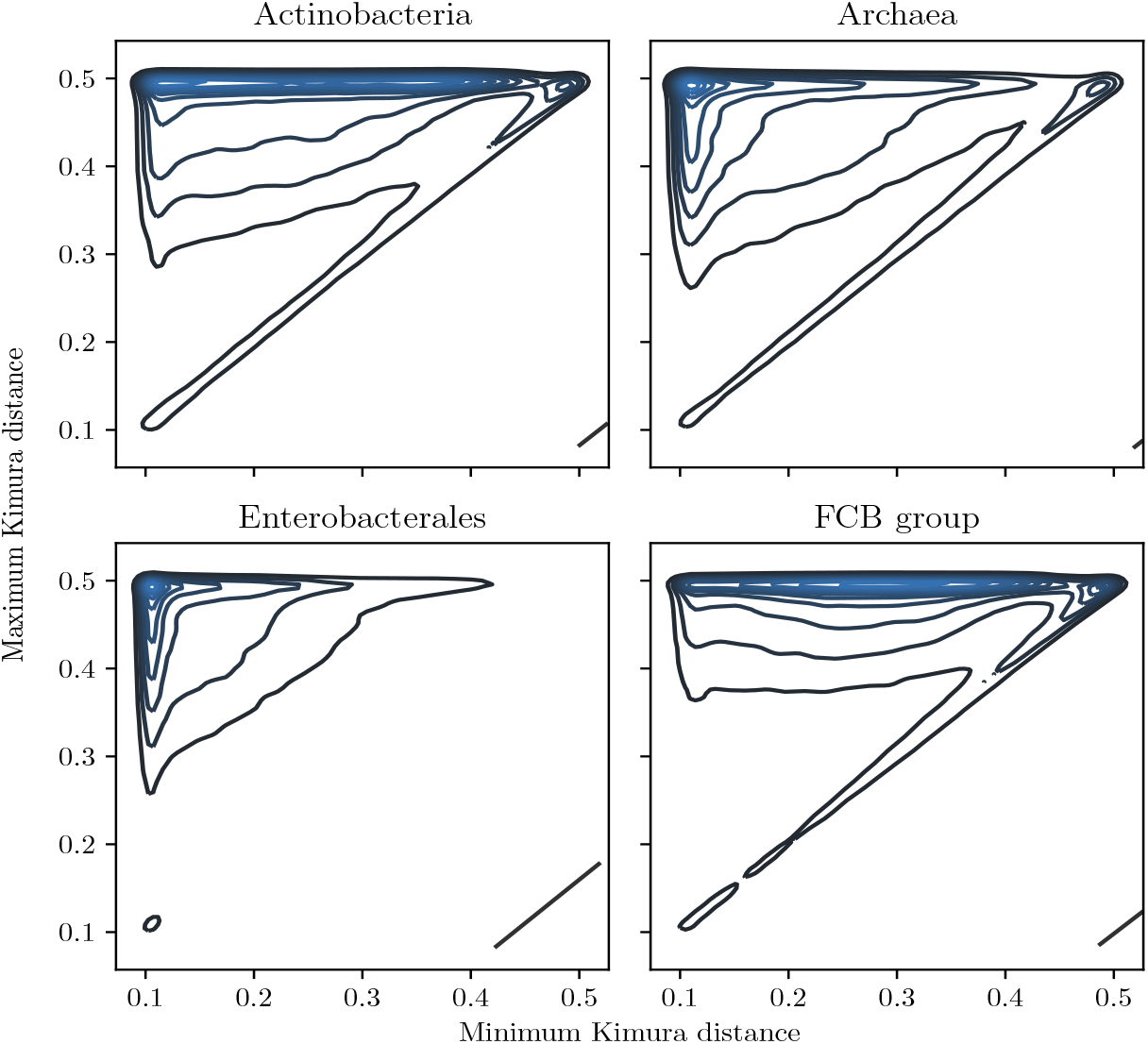
The distribution of (min K(q), max K(q)) vectors defined for queries and their homologs. Data on more than 1,000,000 query genes from 443 genomes (Table 1).

In *Actinobacteria* and *FCB group,* however, large fractions of the query genes had the closest relatives at a rather long distance with the minimum Kimura distance varying from 0.1 to 0.4. Therefore, the average Kimura distance per query was 0.38 for *Actinobacteria* and *FCB group* compared to 0.23 for *Enterobacterales* (Fig. 11). We observed that the homologs of genes of *Enterobacterales* species span uniformly a broad range of the Kimura distances (from the respective query genes). Such distributions produced a robust performance of StartLink as well as high coverage of genes in a query genome by the StartLink predictions.

**Figure 11:**
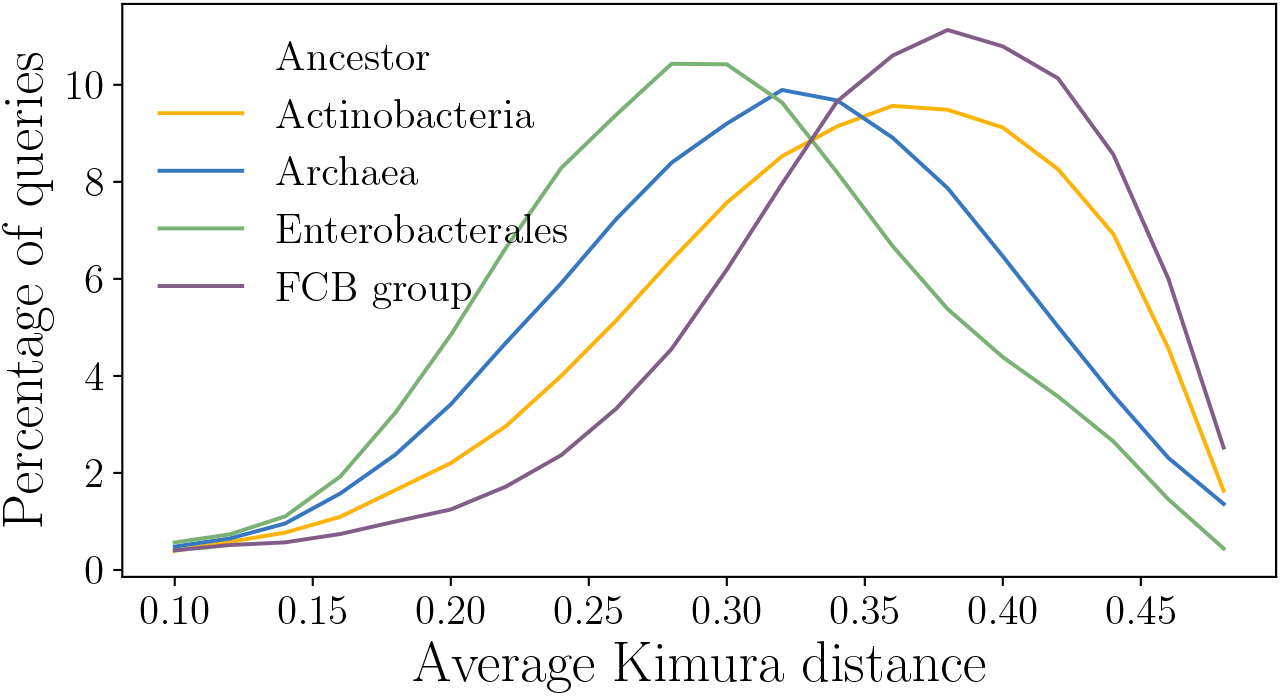
The four clades distributions of the average Kimura distances from a query to the homologs. The y-axis shows the percentage of queries that had a particular average Kimura distance to their homologs.

On the set of genes with verified starts we found StartLink’s and especially StartLink+’s error rates to be uniformly low regardless of the Kimura distance range (see Figs. S2, S3). We also observed that deviations of the StartLink+ predictions from the PGAP annotation were in the same range regardless of variations in min and max values of the range of the Kimura between queries and targets (Fig. S5).

### Variability of the BLAST hits distributions across different clades

Besides the variability in the Kimura distance distributions, the four prokaryotic clades also showed clade-specific variability among query genes with respect to the numbers of homologs detected in similarity searches. A distribution of the number of BLASTp hits (prior to any filtering) in each of the four clades is shown in Fig. 12a, while the percentages of query genes (per genome) that had at least *N* BLASTp hits, where *N* varies from 0 to 5,000 hits is shown in Fig. 12b.

**Figure. 12:**
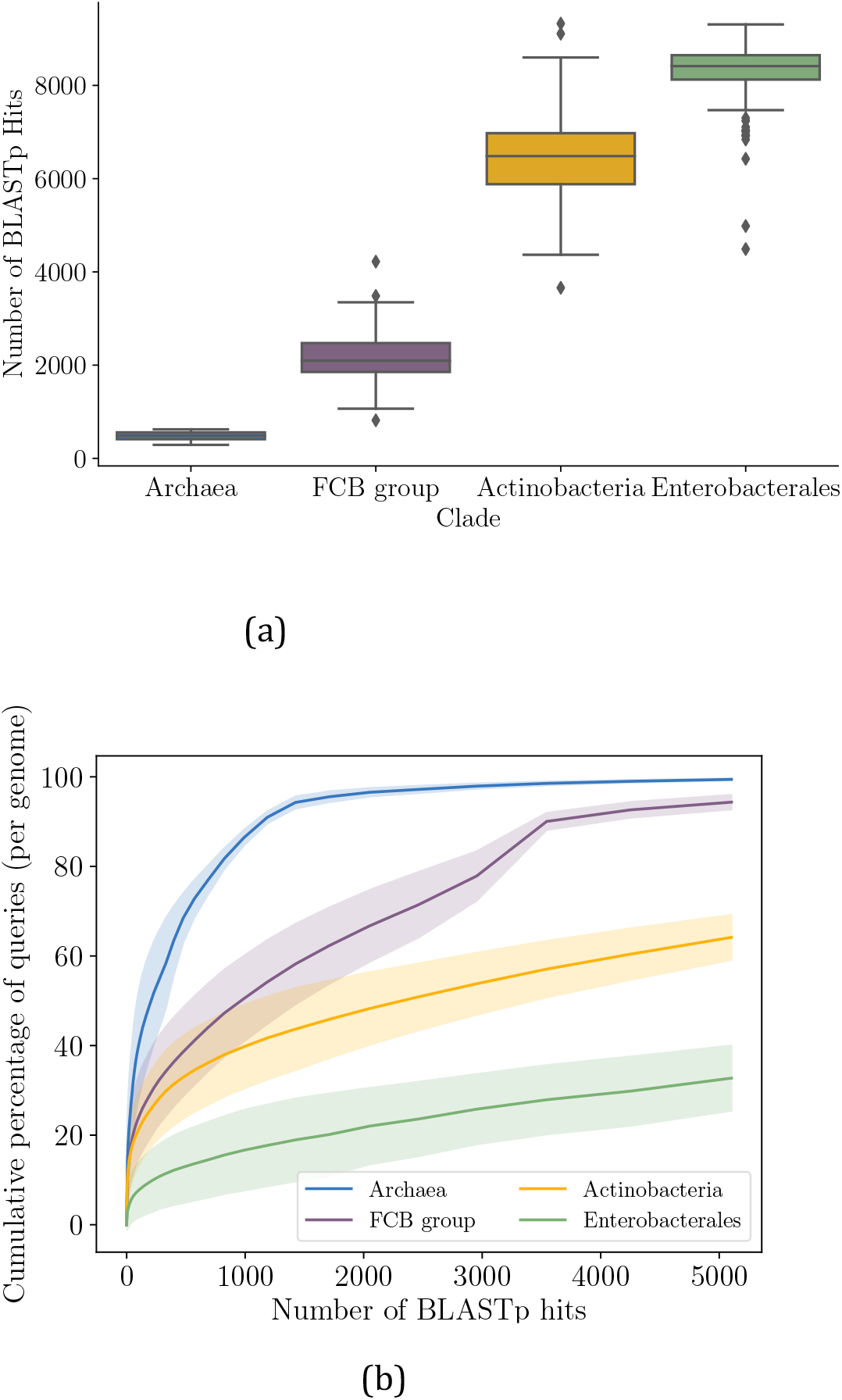
Distribution of raw BLASTp hits per queries in different clades (Table 1). (a) Box plots of the raw numbers of BLASTp hits per query (b) The cumulative percentage of queries with at most N BLASTp hits. The shaded bands show the standard deviations (per clade) across query genomes.

Naturally, the number of hits per query was largely proportional to the number of genomes within a clade (Table 1, Fig. 12a). On the other hand, the cumulative distributions (Fig. 12b) rose very quickly and plateau early on, first for *Archaea* (1,125 genomes) and then the *FCB group* (3,306 genomes). In *Enterobacterales* (6,311 genomes) and *Actinobacteria* (8,097) the cumulative distributions grew much more slowly. Still *Actinobacteria’s* distribution (which has *more* genomes) grew significantly faster than *Enterobacterales*. For example, the likelihood that a query in *Enterobacterales* got *at least* 1,000 BLAST hits was ≈ 83%, compared to only 60% in *Actinobacteria*.

### Visualization of the StartLink data analysis

The multiple sequence alignments used for the StartLink inference could be of interest for visual inspection of the pattern of conservation. For example, an MSA made for a gene adhE1 Rv0162c in *M. tuberculosis* showed a case where StartLink+ prediction was different from the annotated gene start (Fig. 13).

**Figure. 13:**
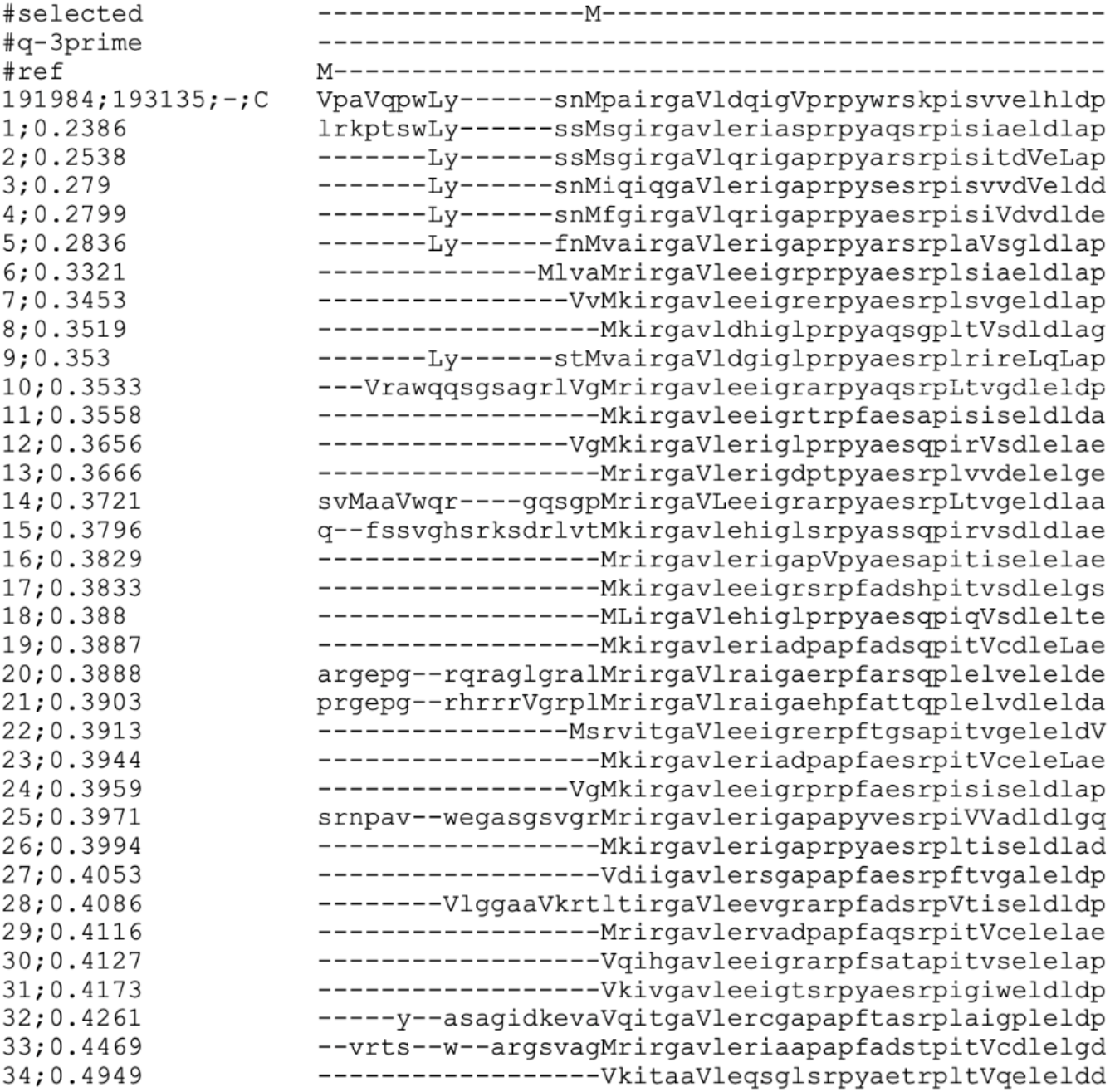
MSA for gene adhE1 Rv0162c in M. tuberculosis, the Actinobacteria clade.

The top amino acid sequence (line 4) is the translated query sequence, followed by the sequences of selected homologs. Capital M, V, and L letters represent Methionine, Valine, and Leucine coded by ATG, GTG, and TTG respectively. Lowercase *v* and *l* represent Valine and Leucine coded by non-GTG or non-TTG codons, respectively.

Annotated start of this gene (“#ref”) was the GTG-coded Valine, while StartLink+ predicted as start the downstream Methionine (“#selected”). We see that StartLink+’s prediction had a high conservation of both the gene-start and the immediate downstream region. Conversely, the annotated start was positioned in a highly non-conserved upstream region. (more MSA examples are shown in Supplementary Note 7).

## Discussion

### Role of the StartLink algorithm

StartLink is a novel algorithm that infers gene start position from analysis of patterns of conservation in syntenic nucleotide sequences. The task of inference complicated by the necessity to select the reference sequences that have to be evolutionarily not too close to as well as not too distant from the sequence of interest (the query). Multiple alignment of sequences selected around the presumable starts of the homologous genes (the query and targets of the BLAST search) is expected to exhibit changes of the positional conservation pattern upon crossing the position of start from to intergenic regions to the genes or from gene in one reading frame to a gene in another frame in the same strains (a gene overlap).

While already existing annotations of syntenic sequences may be useful, the automatic transfer of gene start annotation may propagate errors made in annotation of the earlier sequenced genomes.

Current similarity-based approaches often rely on protein databases, aligning the full length of (or a downstream fragment within) these proteins to detect protein conservation. Obvious drawback of such approach is the absence of a search for conservation in longer sequences and risk of propagation errors made in annotation of the earlier sequenced genomes (Wall et al. 2011).

### Comparing StartLink+ with Prodigal and PGAP

StartLink+ has demonstrated a uniformly low error rate in gene start prediction on the sets of genes with experimentally verified starts. On the other hand, noticeable differences were observed between PGAP and StartLink+ start predictions. These differences were not uniform among the outcomes of the A, B and C algorithmic steps of StartLink+ (Fig. 7). Specifically, we observed fewer differences with predictions made at step A which is logical since if a single candidate start exists upstream of a region determined to be protein-coding, then this candidate is the true start.

The difference across clades (Fig. 5) may be due to multiple factors, such as the different distributions of available orthologs across Kimura distances, as shown in Fig. 10. This has to be accounted for in any comparative approach; we show that StartLink+ performs reliably across the Kimura distances (Supplementary Note 2). Similarly, StartLink’s selection of target sequences is gene-specific instead of genome-specific, to account for genes that evolve at different speeds. These differences lead to different sets of target sequences between PGAP and StartLink, which has an effect on the start-selection algorithm.

The behavior across GC is similar when comparing StartLink+ to PGAP and to Prodigal. However, while the difference between StartLink+ and Prodigal was the highest for *Actinobacteria* and increased with GC, it decreased after the 67% GC (Fig. 5). Also, a larger difference in starts with Prodigal was observed for the FCB group.

Notably, on the set of 5,488 representative genomes the average gene-start differences between StartLink+ and annotation showed the similar increase with increase of the GC content (Fig. 1) as the ones observed upon comparison of the gene finding algorithms on a smaller set of genomes from four clades (Fig. 5).

### Coverage of the whole complement of genes by predictions made by StartLink and StartLink+

StartLink made on average 85% coverage over the gene complement of a prokaryotic genome (Fig. 14a), with the *Enterobacterales* average, 92% per genome, significantly higher than 80-83% average observed in the remaining three clades.

**Figure. 14:**
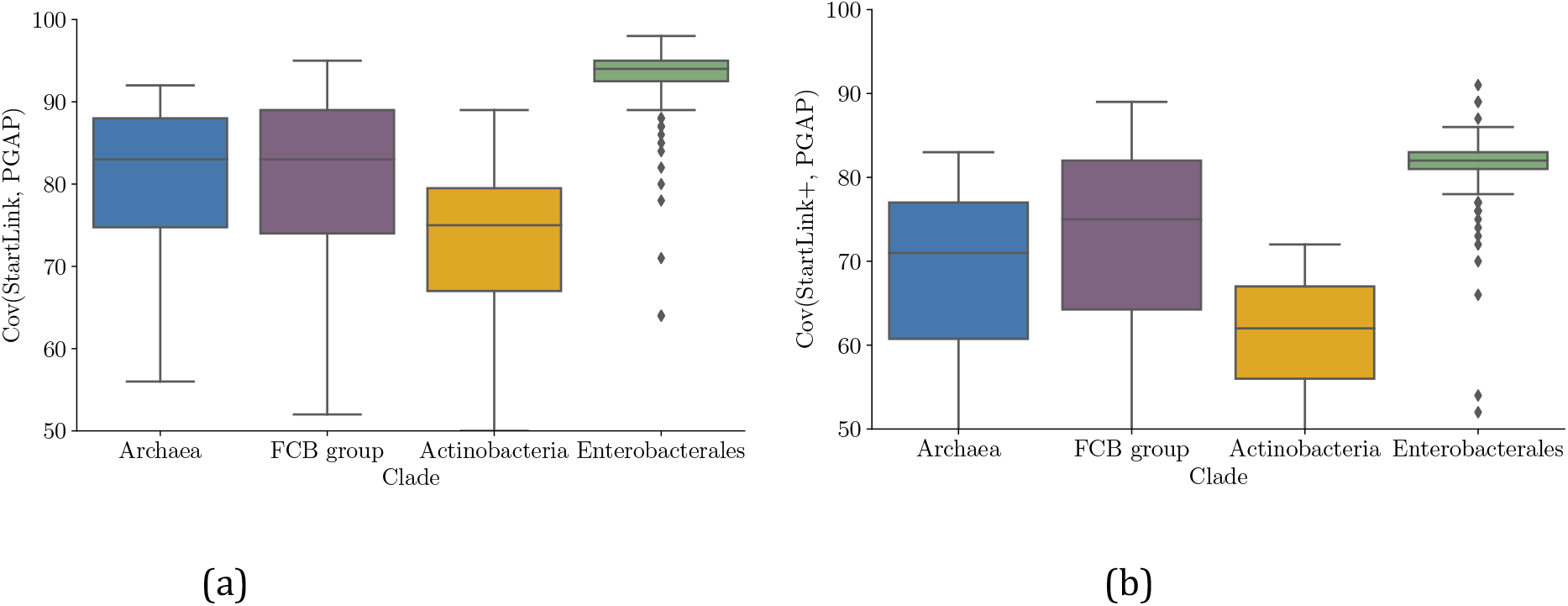
Coverage rates of (a) StartLink and (b) StartLink+ for the four clades. The analysis was done on the same data as for Fig. 5.

The StartLink coverage per genome was expected to depend on the phylogenetic position of the species and the pattern of sampling of close species for whole genome sequencing. Specifically, the percentage of genes where there are no raw BLASTp hits might provide an upper bound for the possible coverage. The cumulative percentage of queries (per genome) that have at most *n* BLASTp hits, where *n* ∈ [0, 40] was shown in Fig. 15. We were interested in queries with fewer BLAST hits than the majority of the genes in a genome (which could be thousands of hits, Fig. 12). Interestingly we saw that on average, 10% of *Archaea*, 12% of *Actinobacteria*, and 12% of *FCB group* genes (per genome) had *fewer* than 10 BLASTp hits, while only 3% of *Enterobacterales* genes had fewer than 10 hits. Moreover, these hits might not land within the desired Kimura distance intervals to the query and to each other. In comparison to the coverage rates of StartLink for each of the clades, we saw that a large percentage of the loss of coverage could be traced back to the low number of hits.

**Figure 15:**
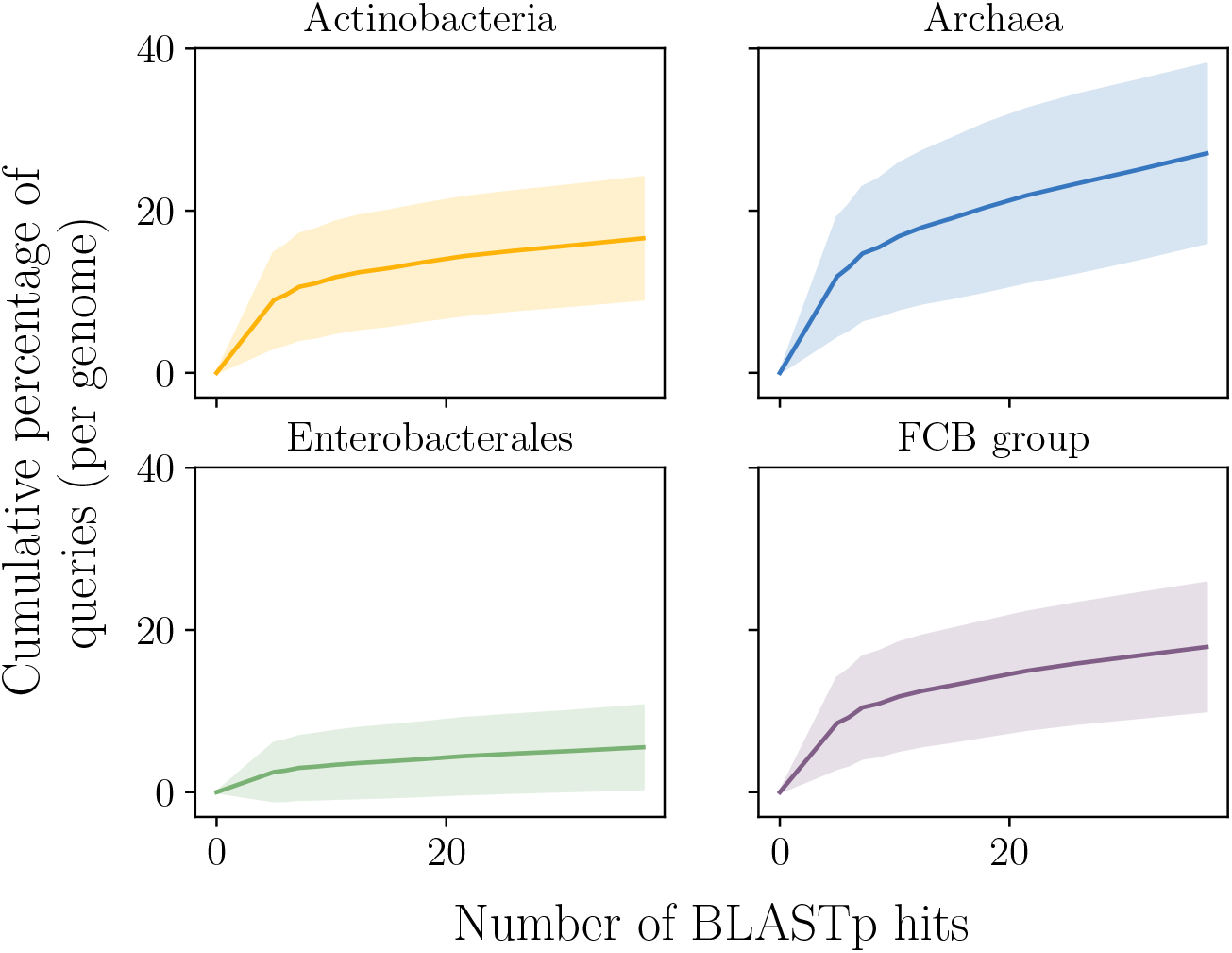
The cumulative distribution of BLASTp hits (<40) per query in a genome, shown for the four clades. This is a zoomed in version of Fig. 12.

The average coverage rate of StartLink+ per genome was 73%; also, the differences between the average coverage per clade became larger (Fig.14b). For instance, *Actinobacteria*’s coverage dropped by 16% from StartLink to StartLink+ (the largest drop across the clades), indicating the increased difference in gene start predictions between GeneMarkS-2 and StartLink.

### Differences in numbers of selected homologs

For the sake of reducing the running time of StartLink, we limited the maximum number of allowed targets used in the MSA (currently *N* = 50). Thus we selected at most *N* sequences from the set of BLASTp hits and continued with further filtering within the MSA (Supplementary note 6).

As such, the effective number of targets per query at the end of a StartLink run can be less than *N*. The average number of targets per query after a full StartLink run can differ significantly between clades (Fig. 16), especially when comparing *Archaea* and *Enterobacterales*.

**Figure 16:**
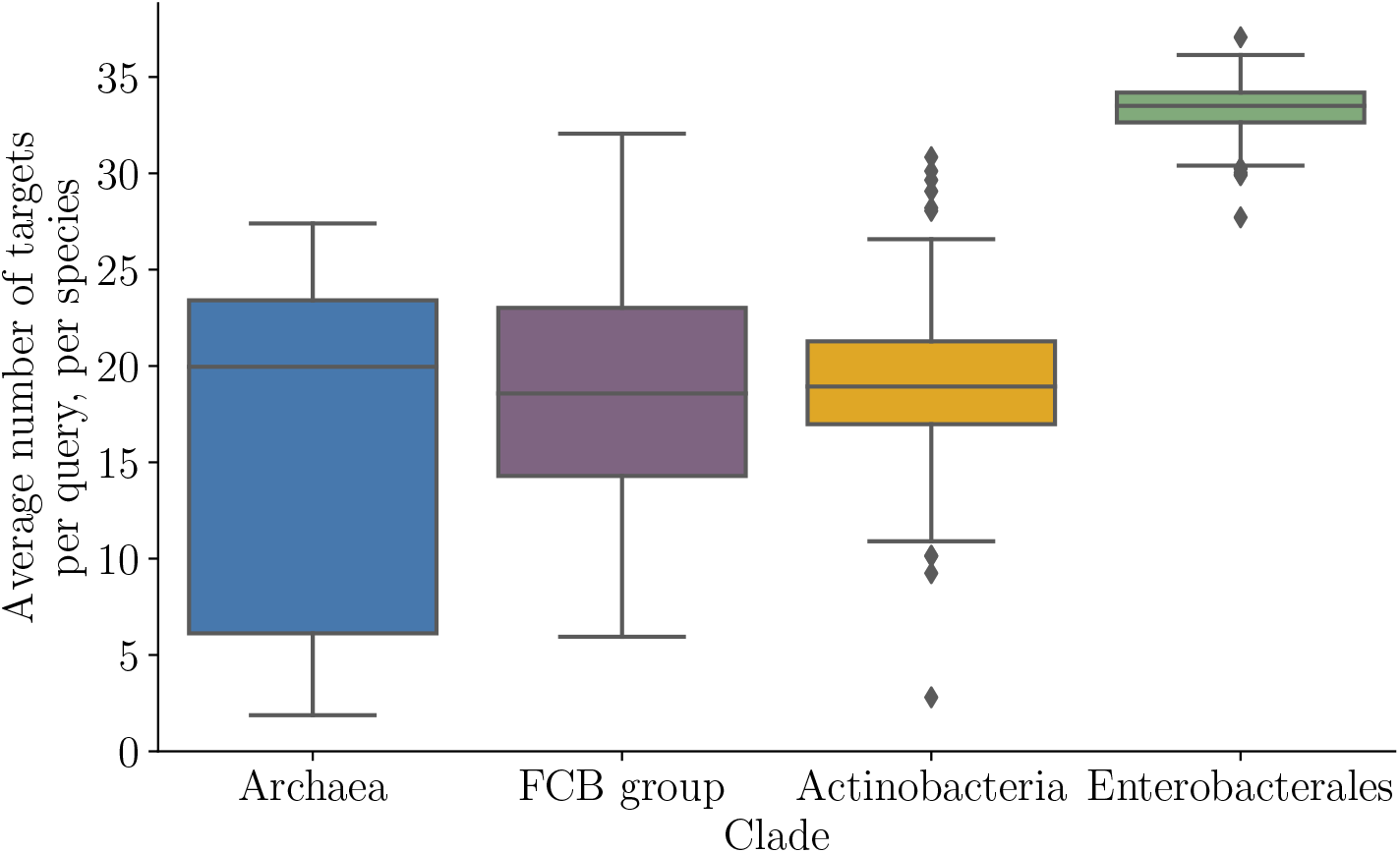
The average number of targets per query in the StartLink runs. The average was computed per genome and shown for each of the four clades (in the whole set of 443 query genomes).

A possible reason why *Enterobacterales* had a high average number of targets is the large spread of the Kimura distances compared to other clades (see Fig. 10). On the other side, the spread of averages within *Archaea* was non-uniform, reaching as low as 10 targets per query on average. This was partly due to the sparseness of species in the phylogenetic space of known archaea and small number of sequences genomes in this clade, making it less likely that we find enough sequences within the right Kimura range.

Nonetheless, we should note that for the set of genes with verified starts, the observed differences in the number of targets per query did not translate into a difference in the StartLink accuracy. For example, when StartLink was run with *N* = 50, both archaea (*H. salinarum* and *N. pharaonis*) end up with on average 20 targets per query, compared to *E. coli* 40 targets per query. However, for *H. salinarum* and *N. pharaonis* we observed an error rates of 3% and 2% respectively, while the *E. coli* error rate was 5%.

To assess the performance of StartLink on *Archaea* with low target-per-query averages we decreased *N* to 20. This change produced 10 to 15 targets per query for both *archaea* species. Still, we saw just a slight increase in error rate for *H. salinarum* (by 0.6%) and a decrease in error rate for *N. pharaonis* by 0.7%. For all the sets of genes with verified starts, we saw 0.5% change in error rate (on average per genome) when *N* was shifted. This result demonstrated that StartLink was stable with respect to the change of *N*, with the minor changes in error rates possibly caused by random selection of target sequences.

## Conclusions

Given that existing computational tools for *ab initio* gene finding in prokaryotic genomes could differ by 15-25% in gene start predictions despite seemingly highly accurate predictions on limited in size sets of genes with verified starts we turned to development of a gene finder supported by evolutionary conservation patterns in known genomic and protein sequences. We introduced StartLink, a comparative genomics based algorithm and tool for gene start prediction working independently from an *ab initio* methods.

Next, we introduced StartLink+, integrating GeneMarkS-2 and StartLink, and have shown that it delivers low error rates in gene-start predictions (~1%) along with a sufficiently high coverage of genes per genome (~73% on average). The gene starts predicted by StartLink+ besides i/ the significance for improving current genome annotations ii/ the role for more accurate learning sequence patterns around gene-starts in finer details, also carry significant potential for iii/ studies of evolution of regulatory sequences near gene starts controlling diverse gene expression mechanisms.

## Supporting information

Full set of Supplementary materials

## Availability

The StartLink+ code and instructions on how to access the datasets needed to reproduce the results presented in this paper are available at the GitHub page: https://github.gatech.edu/GeneMark/startlink

## Funding

This work was supported in part by the National Institutes of Health (NIH) [GM128145 to M.B.]. Funding for open access charge: National Institutes of Health [GM128145].

## Conflict of interest statement

None declared.

## Notes

https://github.gatech.edu/GeneMark/startlink

### Competing Interest Statement

The authors have declared no competing interest.

