## Supplementary material for "StartLink+: Prediction of Gene Starts in Prokaryotic Genomes by an Algorithm Integrating Independent Sources of Evidence": Full set of Supplementary materials

### Note 1: Accuracy of combined predictors

In pattern recognition, adding up an independent and informed method of analysis and prediction leads to significant increase in accuracy. This principle is often applied in order to corroborate an evidence for an inference from data with incomplete information. The task of gene-start prediction is exactly one where such a principle could be applied.

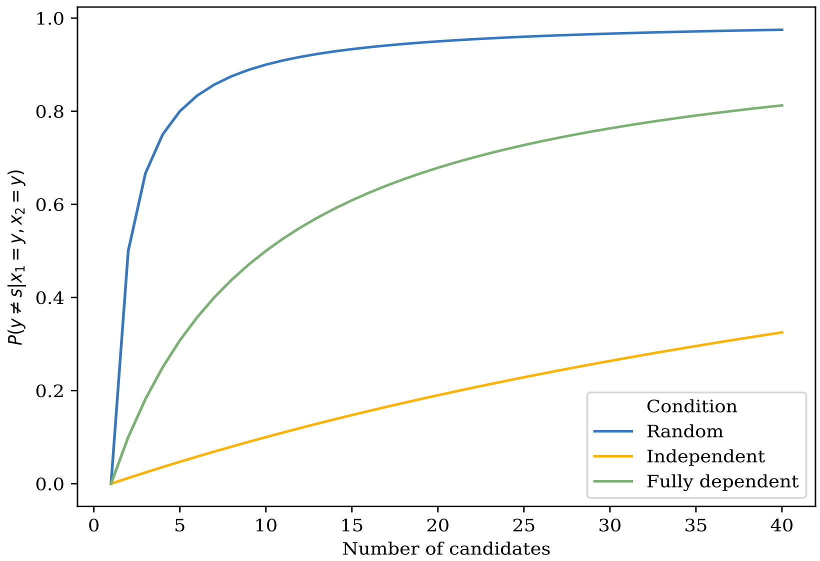


*Fig. S1: The probability that an identical selection made by two algorithms $A_1$ and $A_2$ is incorrect, as a function of the number of candidates for the selection. The plots are shown for the three dependency conditions between $A_1$ and $A_2$. Err (Ai = 0.1), except for the random case where the error is the fraction of false candidates.*

Suppose we have two algorithms, $A_1$ and $A_2$, whose gene-start predictions $x_1$ and $x_2$, respectively, have average error rates Err$(A_1)$, Err$(A_2)$. For a gene with candidate starts $C = \{c_1, \dots, c_{|C|}\}$ and a **true start** $s \in C$, we ask the following question: If algorithms $A_1$ and $A_2$ predict the same start $y$ (i.e. $x_1 = x_2 = y$), what is the probability that $y$ is the true start (i.e. $y = s$)?

$$p(y = s | x_1 = y, x_2 = y) = ? \quad (1)$$

We have:

$$\begin{aligned} p(x_i = y | y = s) &= 1 - Err(A_i) = \text{Acc}(A_i) \\ p(x_i = y | y \neq s) &= \text{Err}(A_i) \end{aligned} \quad (2)$$

Then we consider the cases when $A_1$ and $A_2$ predict the same start

1. completely independent
2. completely dependent (e.g., the same algorithm),
3. select a start at random among several candidates with equal probability

   Deriving Eq 1 for independence ($P_I$), and full dependence ($P_D$), randomness ($P_R$) conditions (see below for the proof), we get

$$\begin{aligned} P_I &= \frac{1}{1 + (|C| - 1)\frac{\text{Err}(A_2)\text{Err}(A_1)}{\text{Acc}(A_2)\text{Acc}(A_1)}} \\ P_D &= \frac{1}{1 + (|C| - 1)\frac{\text{Err}(A_2)}{\text{Acc}(A_2)}} \\ P_R &= \frac{1}{|C|} \end{aligned} \quad (3)$$

Since $p(x_i = y | y \neq s) = 1 - p(x_i = y | y = s)$, the probability of the identical prediction being false increases (Fig. S1) as the number of options to select from increases. More importantly, it shows the effect of adding a second, independent prediction on the overall error rate, indicating that the largest improvement can be gained by maximizing independence between the algorithms.

Note that the real error rates of gene-start prediction are difficult to model theoretically, and the above consideration makes simplifying assumptions (e.g., on the uniformity of selecting any of the incorrect starts when a wrong prediction is made).

#### Proof of Equation 3

The derivations of Eq 2 under the three conditions of randomness, independence, and full dependence, is as follows. The first expansion (irrespective of condition) is

$$
\begin{aligned}
&p(y = s|x_1 = y, x_2 = y) \\
&= \frac{p(x_1 = y, x_2 = y, y = s)}{p(x_1 = y, x_2 = y)} \\
&= \frac{p(x_1 = y, x_2 = y, y = s)}{p(x_1 = y, x_2 = y, y = s) + p(x_1 = y, x_2 = y, y \neq s)} \\
&= \frac{1}{1 + \frac{p(x_1 = y, x_2 = y, y \neq s)}{p(x_1 = y, x_2 = y, y = s)}} \\
&= \frac{1}{1 + \frac{p(y \neq s)\, p(x_2 = y|y \neq s)\, p(x_1 = y|x_2 = y, y \neq s)}{p(y = s)\, p(x_2 = y|y = s)\, p(x_1 = y|x_2 = y, y = s)}} \\
&= \frac{1}{1 + \frac{p(y \neq s)\, \mathrm{Err}(A_2)\, p(x_1 = y|x_2 = y, y \neq s)}{p(y = s)\, \mathrm{Acc}(A_2)\, p(x_1 = y|x_2 = y, y = s)}}
\end{aligned} \tag{4}
$$

Note that assuming uniform priors, we have

$$
\begin{aligned}
P(y = s) &= \frac{1}{|C|} \\
P(y \neq s) &= \frac{|C| - 1}{|C|}
\end{aligned} \tag{5}
$$

Specifically, the prior over $p(y \neq s)$ assumes that when an algorithm makes a wrong prediction, it is equally likely to choose any of the false candidates. In reality, this may or may not be true, especially if algorithms tend to choose false starts closer to the true start. This is important especially under conditions where algorithms are somewhat dependent. However, given that we have no knowledge of $A_1$ and $A_2$, the most straightforward approach is to assume an uninformed prior. Therefore, we get

$$
\begin{aligned}
&p(y = s|x_1 = y, x_2 = y) \\
&= \frac{1}{1 + (|C| - 1)\frac{\mathrm{Err}(A_2)\, p(x_1 = y|x_2 = y, y \neq s)}{\mathrm{Acc}(A_2)\, p(x_1 = y|x_2 = y, y = s)}}
\end{aligned} \tag{6}
$$

**Case 1:** Algorithms $A_1$ and $A_2$ are uniform, random selectors.
Since $A_1$ and $A_2$ are completely random, then they are (by definition) independent. This implies that

$$
\begin{aligned}
p(x_1 = y|x_2 = y, y = s) &= p(x_1 = y|y = s) = \frac{1}{|C|} \\
p(x_1 = y|x_2 = y, y \neq s) &= p(x_1 = y|y \neq s) = \frac{|C| - 1}{|C|}
\end{aligned} \tag{7}
$$

Therefore, we have

$$
\boxed{p(y = s|x_1 = y, x_2 = y) = \frac{1}{|C|}} \tag{8}
$$

**Case 2:** Algorithms $A_1$ and $A_2$ are completely **independent**
Again, since $A_1$ and $A_2$ are independent, we have

$$p(x_1 = y | x_2 = y, y = s) \quad = p(x_1 = y | y = s) = \text{Acc}(A_1) \tag{9}$$
$$p(x_1 = y | x_2 = y, y \neq s) \quad = p(x_1 = y | y \neq s) = \text{Err}(A_1)$$

Therefore, we have

$$\boxed{p(y = s | x_1 = y, x_2 = y) = \frac{1}{1 + (|C| - 1)\frac{\text{Err}(A_2)\text{Err}(A_1)}{\text{Acc}(A_2)\text{Acc}(A_1)}}} \tag{10}$$

As an example, the values for equation 10 for C = 2, 3, 4, and 5 are 0.99, 0.98, 0.96, 0.95, 0.94

**Case 3:** Algorithms $A_1$ and $A_2$ are completely **dependent**
Again, since $A_1$ and $A_2$ are independent, we have

$$p(x_1 = y | x_2 = y, y = s) \quad = 1 \tag{11}$$
$$p(x_1 = y | x_2 = y, y \neq s) \quad = 1$$

Therefore, we have

$$\boxed{p(y = s | x_1 = y, x_2 = y) = \frac{1}{1 + (|C| - 1)\frac{\text{Err}(A_2)}{\text{Acc}(A_2)}}} \tag{12}$$

### Note 2: Effects of the variations in distributions of the Kimura distance on the StartLink performance

StartLink uses the Kimura distances as a metric to filter out sequences of too close or too distant relatives. We tested several alternative evolutionary distance metrics, including (non-) synonymous substitution rates, amino acid identity, etc. but they either performed equally well or slightly worse at providing StartLink with a good set of orthologous sequences (data not shown). Overall, we found that *these* metrics provide a way to remove sequences of evolutionarily very close/distant relatives, but that additional filtering (such as analyzing the gaps in the MSA incurred by each query, as described in the main text) might still be needed.

The effect of varying the maximum Kimura threshold from 0.2 to 0.8, while fixing the minimum threshold to 0.1, is shown in Fig. S2. StartLink's sensitivity is robust with respect to increase of the maximum Kimura distance. On the other hand, the coverage rate fluctuates. This occurs because when more sequences with larger Kimura distance from query are included in the MSA (and thus, more mutations), the gene start becomes more difficult to infer.

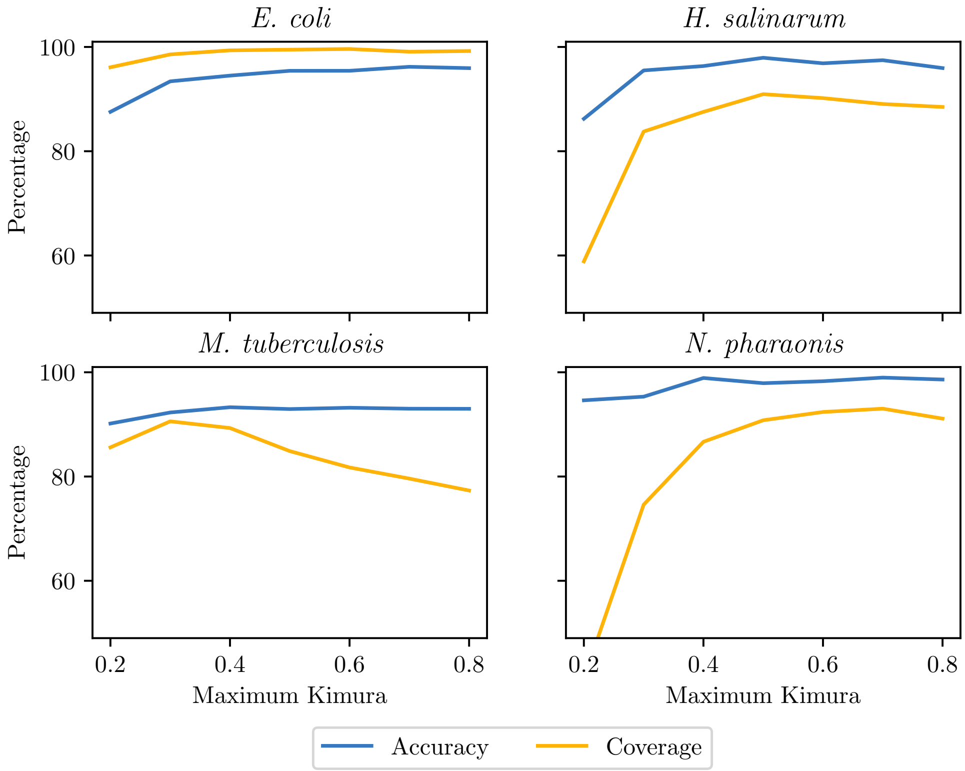


*Fig. S2: The effect of changing the maximum Kimura threshold on StartLink's sensitivity and coverage rates. The minimum Kimura threshold is fixed to 0.1, and* $x \in \{0.2, 0.3, \ldots, 0.8\}$.

Specifically, *N. pharaonis* and *H. salinarum*'s coverage rates suffer when the maximum Kimura threshold is below 0.4. Note, however, that the number of available genomes of Archaea (used as a reference clade) is quite small, especially when compared to numbers in *Enterobacterales* and *Actinobacteria*. This means that, in general, there are fewer close sequences that could directly impact the coverage rate.

On the other hand, *M. tuberculosis*'s coverage rate decreases as the maximum threshold increases past 0.4. A direct inspection of the MSAs showed that they appear to have more mutations around the gene-start region than, for example, in *E. coli* for the same range of Kimura distances. It is unclear whether this is due to a sampling bias of genomes in the database or a biological feature. That said, StartLink+'s sensitivity rate remains constant, meaning that for such alignments, it does not make a prediction.

Similarly, Fig. S3 fixes the maximum Kimura threshold to 0.5 and varies the minimum threshold from 0.001 to 0.4. Overall, this has a lower effect on the coverage and sensitivity rates than in the previous case. We see an initial drop in the case of *E. coli* when the minimum threshold is close to 0.001. In this range, we observed more orthologs with high nucleotide-level identity rate (to the query gene) than in the other three genomes. It is again unclear whether this stems from a genome sampling bias in the database or from some underlying patterns of genome evolution, but the overall reduction in sensitivity is still small.

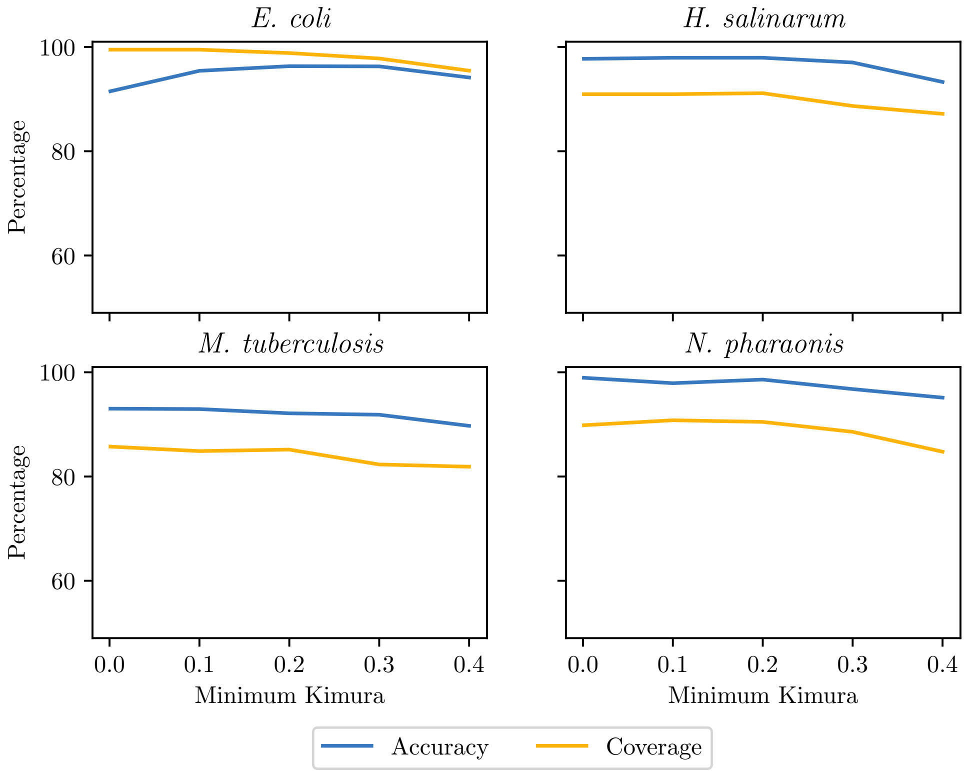


*Fig. S3: The effect of changing the minimum Kimura threshold on StartLink's sensitivity and coverage rates. The maximum Kimura threshold is fixed to 0.5, and* $x \in \{0.001, 0.1, 0.2, 0.3, 0.4\}$.

It is important to note that for all the above experiments, the sensitivity of StartLink+ remains consistently high and largely unaffected, while the coverage rate mirrors that of StartLink since it is directly dependent on it (data not shown). In other words, our selection of the Kimura distances interval does not need to be fine-tuned beyond what has been done. This is fortunate, given that we do not want to overfit the small size ground-truth data available to us.

Finally, we tested StartLink on small intervals of the Kimura distance, specifically $[0.001, 0.1]$, $[0.1, 0.2]$, … $[0.7, 0.8]$. This forced StartLink to operate exclusively in regions of evolutionarily very close and very distant orthologs, and allowed us to see a more detailed view of the effect of the Kimura distance on the algorithm. Fig. S4 shows the effect of these ranges on the sensitivity and specificity.

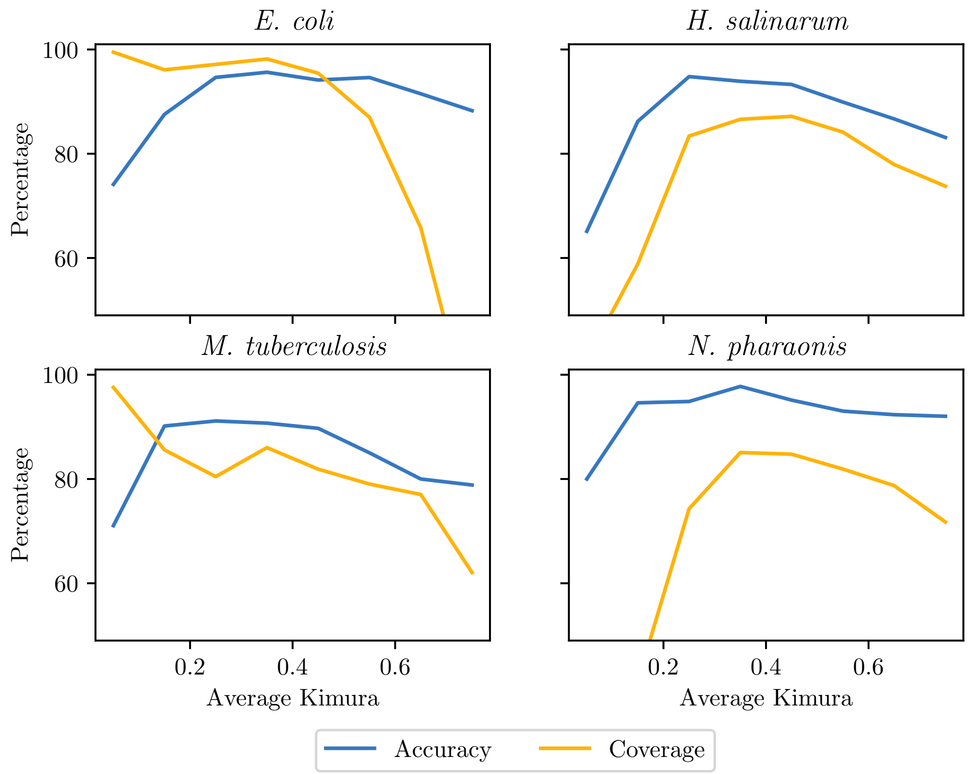


*Fig. S4: The performance of StartLink on small intervals of the Kimura distance ranges:* $[0.001,0.1]$, $[0.1,0.2]$, $[0.2,0.3]$ … $[0.7,0.8]$. *The x-axis shows the mean Kimura distance; e.g., for range* $[a, b]$*, the average is* $\frac{b+a}{2}$.

The effect here is more significant, especially at the blocks at the edges of the whole range, i.e. $[0.001,0.1]$ and$[0.7,0.8]$. The sensitivity is still rather stable, with the biggest impact being in the range $[0.001,0.1]$. The coverage, however, is changing significantly. First, the coverage behaves differently in *E. coli* and *M. tuberculosis* compared to the two archaeal species. Specifically, it starts out high in both *E. coli* and *M. tuberculosis*, and then drops down beyond the $[0.4,0.5]$ range. The drop is especially strong in *E. coli*, where StartLink struggles to make predictions for ranges [0.6, 0.7] and above (although still maintaining a high and stable sensitivity).

On the contrary, the coverage is low for *H. salinarum* and (especially) *N. pharaonis* for the [0.001, 0.1] and [0.1, 0.2] ranges. It then rises quickly and stabilizes, before dropping a little in the higher ranges. The reasons for this pattern are, in part, due to just over 1,000 genomes of Archaea in the entire clade, compared to over 6,311 and 8,097 genomes for *Enterobacterales* and *Actinobacteria*, respectively, with much lower nodes in the taxonomy tree. In fact, *E. coli* and *M. tuberculosis* each have $1{,}000+$ sequenced genomes of the strains within their own species.

Therefore, the observed difference in patterns is likely to only be a result of the undersampling of the microbial species, and may disappear once the database is populated more uniformly.

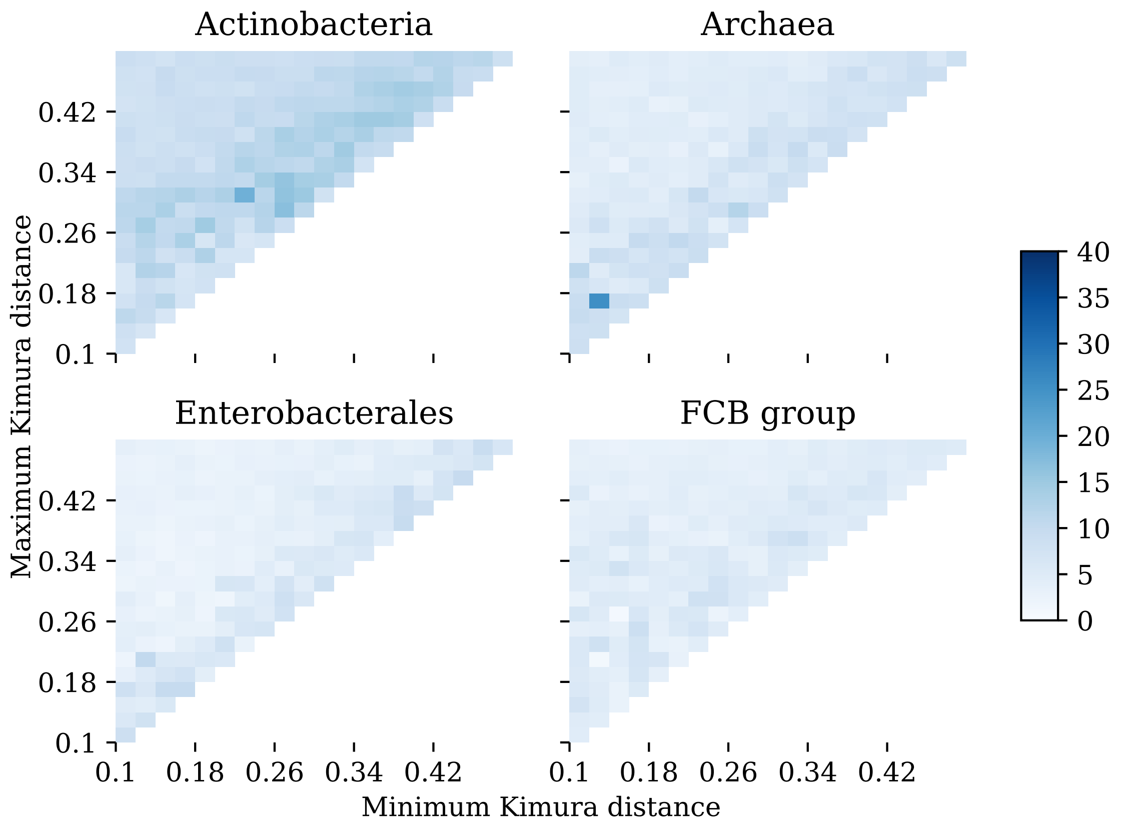


*Fig. S5: The percentage of gene start differences between PGAP and StartLink+, computed as Diff(PGAP, StartLink+), as a function of the minimum and maximum Kimura distances between a query and its targets. The color coding key panel shows the correspondence of color and the percentage of gene starts differences. This analysis was done for 443 query genomes.*

We observed that deviations of the StartLink+ predictions from the PGAP annotation were in the same range regardless of variations in min and max values of the range of the Kimura between queries and targets Fig. S5.

### Note 3: Distributions of the MSA scores computed around gene-starts

The StartLink algorithm relies on metrics, such as a measure of sequence conservation, computed in non-coding and coding regions, thus, around candidate starts. These metrics are computed after removing evolutionarily close and distant sequences, as determined by the Kimura distance, from the set of sequences in the initial MSA.

In this section, we analyze the scores computed by StartLink in different sections of the multiple sequence alignment, from 5' downstream. We run StartLink for genes with experimentally verified starts and build multiple sequence alignments. The goal is to show how the two scores defined in Eq 6 and 7 of the main text behave in different regions of the MSA (specifically, upstream of, at, and downstream of the verified start).

First, we emphasize the following: this analysis is done to show how the algorithm behaves. This means that conditions that StartLink uses, such as only extracting LORFs and filtering by Kimura, are inherently part of the analysis. For example, if a query's verified start is at the LORF 5' end, then the scores for the non-coding region are not computed, StartLink will not search for a gene-start there.

#### Block conservation scores

We start by analyzing the block conservation score $S_{blk}$ defined by Eq 6 of the main text. For each protein MSA, we take blocks (with width 10 *aa*) in four different regions (Fig. S6): 20 *aa* upstream of the verified start (Up-far), just upstream of the verified start (Up-close), just downstream of it (Down-close), and 20 *aa* downstream of the verified start (Down-far).

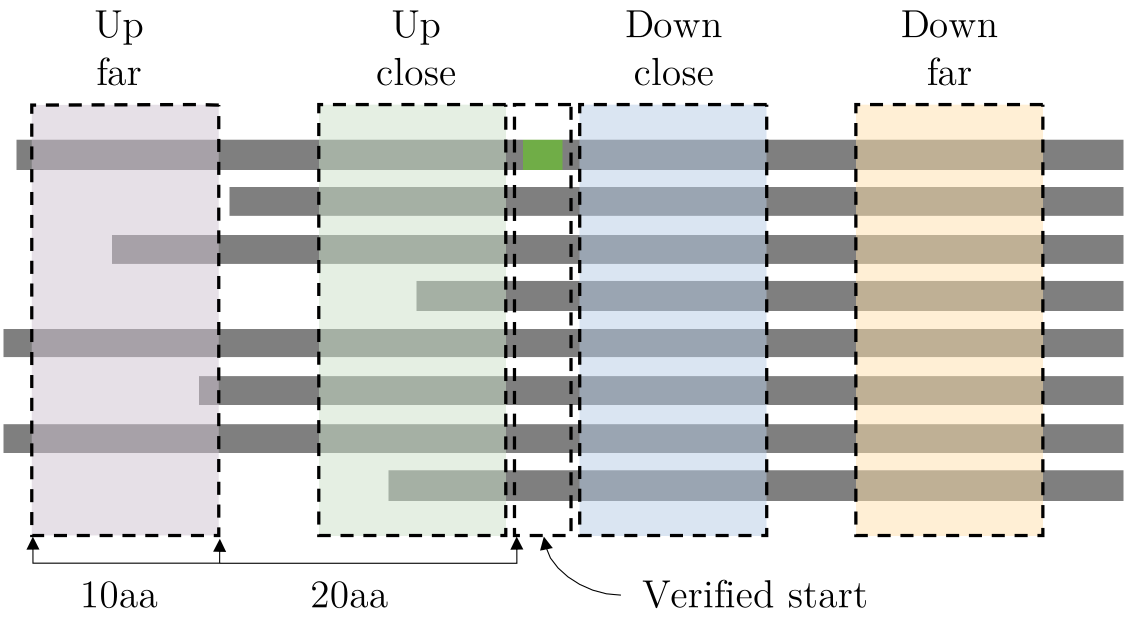


*Fig. S6: Blocks selected for the MSA score analysis.*

Fig. S7 shows the distribution of scores for each block region. As expected, the upstream (non-coding) regions have a much lower conservation score than the downstream (coding) regions, as mutations happen with higher frequency in non-coding regions than in coding regions, Also, given that sequences were extracted as LORFs, this region of the MSA would have a large number of gaps that reduce overall score. This pattern helps StartLink to skip over such regions. Notice that the rules of the StartLink setup (Kimura filtering, LORF extraction, etc...) facilitate the use a simple metric based on identity conservation to skip over non-coding regions, where conserved blocks are not detected.

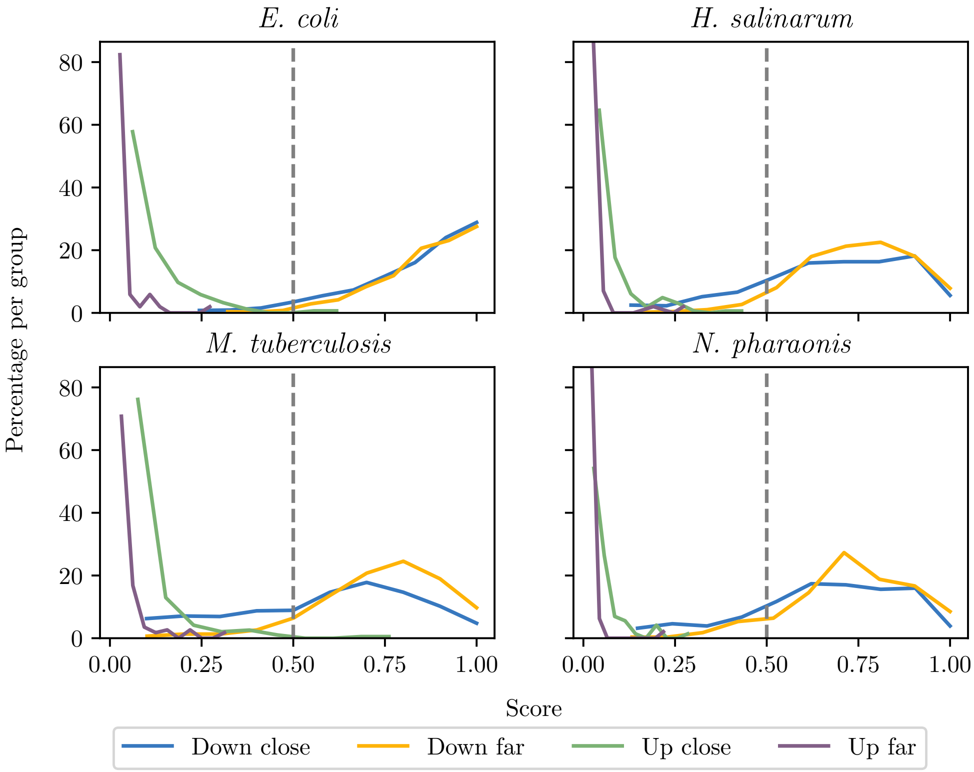


*Fig. S7: Distribution of block conservation scores in regions around verified starts.*

On the opposite end of the protein MSA, the two blocks downstream of the verified start show relatively high conservation scores. Interestingly, the Down-far block has a higher conservation score (on average) than the Down-close one. This is especially apparent in *M. tuberculosis*. This pattern indicates that non-synonymous mutations just downstream of the gene-start may be more likely to occur than in a randomly selected section deeper in a gene. This may be a reason why computation of evolutionary distance metrics on the entire gene may not select an optimal set of sequences for StartLink's gene-start search algorithm.

### Identity score for gene start candidates

Here we analyze the effectiveness of the 5' score (Eq 7 of the main text) for separating true and false starts. Notably, there are two types of false starts.

In an ideal case, the true start will have a high 5' identity score, due to presence of orthologous genes with the same location of the 5' end. The upstream (false) candidates tend to have low conservation scores, due to mutations that occur in non-coding region. On the other hand, the (false) candidates downstream of the verified start can show high identity scores, due to conservation across orthologs. Still, the false starts downstream could be discriminated if codons synonymous to GTG and TTG appear in the alignment to betray absence of selection pressure expected for a start codon. Hence, the penalty for the synonymous codons was added in the equation for $S_{5'}$ (see main text).

Fig. S8 shows how the 5' score separates the annotated (in this case, verified) gene-starts from the false candidates. The gap between the groups is strikingly wide, meaning that the 0.5 threshold is a reasonable, non-overfitting value.

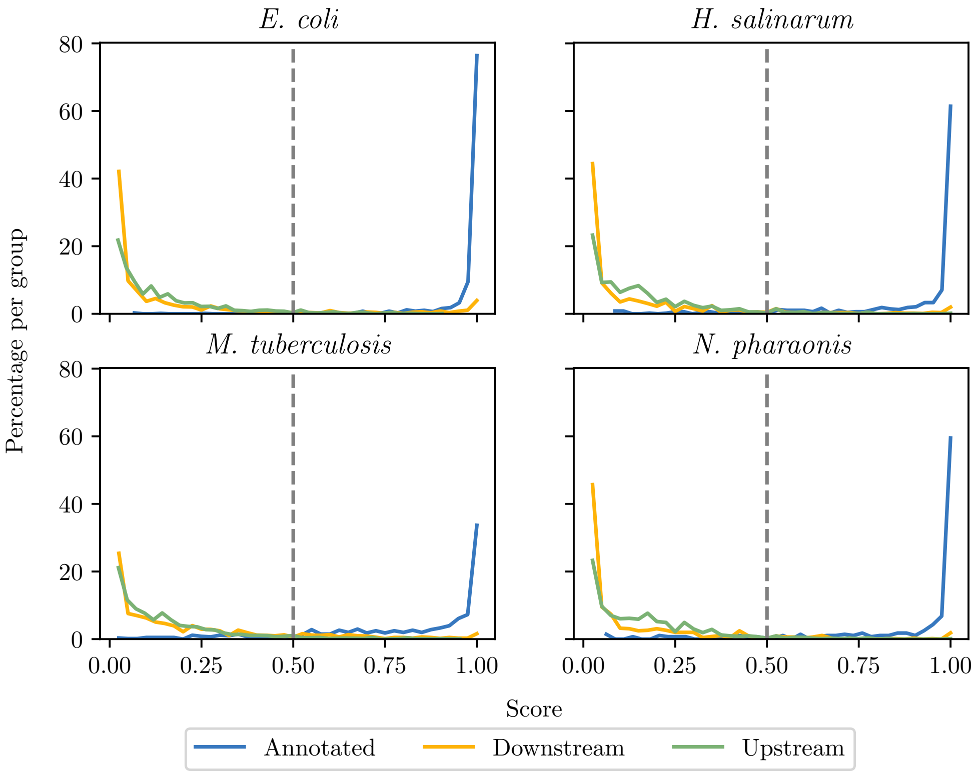


*Fig. S8: Distribution of 5' identity scores for verified starts as well as for upstream and downstream false gene start candidates.*

### Note 4: Distributions of differences in numbers of selected orthologs

The maximum number of allowed targets (currently $N = 50$) is used to limit the number of sequences in MSA. At most $N$ sequences are selected from a large number of BLAST hits and used in the MSA prior to further filtering. The average number of targets per query after a full StartLink run is shown in Fig. 16; it can vary significantly, especially when comparing *Archaea* and *Enterobacterales*.

One reason why *Enterobacterales* has a high average number of targets is the large spread of the Kimura distances compared to other clades (see Fig. 10). On the other side, the spread of Kimura distances within *Archaea* targets has narrow spread, while the average number of targets reaches as low as 10 per query. This is partly due to the small number of genomes in this clade, making it less likely that we find enough sequences within the right Kimura distance range.

Still, we observed on the verified set that the larger final number of targets per query did not translate into a large improvement in performance. For example, for StartLink running with $N = 50$, both archaea (*H. salinarum* and *N. pharaonis*) end up with on average 20 targets per query, compared to *E. coli*'s 40 targets per query. In terms of error rate, however, for *H. salinarum* and *N. pharaonis* we saw the error rate of 3% and 2% respectively, while *E. coli*'s the error rate was 5%.

In another experiment we decreased $N$ to 20 in order to check the performance on Archaea with low target-per-query averages. This change in N resulted in 10 to 15 targets per query for both archaea. However, we saw no real change in error rate; we saw a slight increase in error rate for *H. salinarum* (by 0.6%) and a decrease in error rate for *N. pharaonis* by 0.7%. Moreover, in all verified genomes, when $N$ was shifted we saw a 0.5% (on average per genome) change in error rate, some positive and some negative. This shows that StartLink is stable with respect to $N$, and the minor shifts could be attributed to the underlying randomness of selecting the target sequences.

### Note 5: Gene-start prediction in metagenomes

Standard self-training *ab initio* predictors usually require a sufficiently long sequence (e.g. a sequenced genome) for estimating the models parameters. In metagenomics applications short read assembled sequences normally are lacking full genomic context. Therefore, conventional gene finders such as GeneMarkS-2 cannot run its full cycle with training of the typical model (Lomsadze et al. 2018) due to insufficient training data. Still, GeneMarkS-2 can run a single iteration which is equivalent to running the gene finder with the MetaGeneMark models (Zhu, Lomsadze, and Borodovsky 2010). Such a run relies on pre-built collection of heuristic GC specific models that do not include models for the regulatory motifs in the gene up\stream regions.

Fig. S9 shows results of an experiment made with a goal to show what GeneMarkS-2 accuracy in gene start finding would be on short genome fragments. Since StartLink only requires the gene itself, it's accuracy is stable irrespective of the length of the genomic fragment (as long as the gene itself is contained within the sequenced fragment).

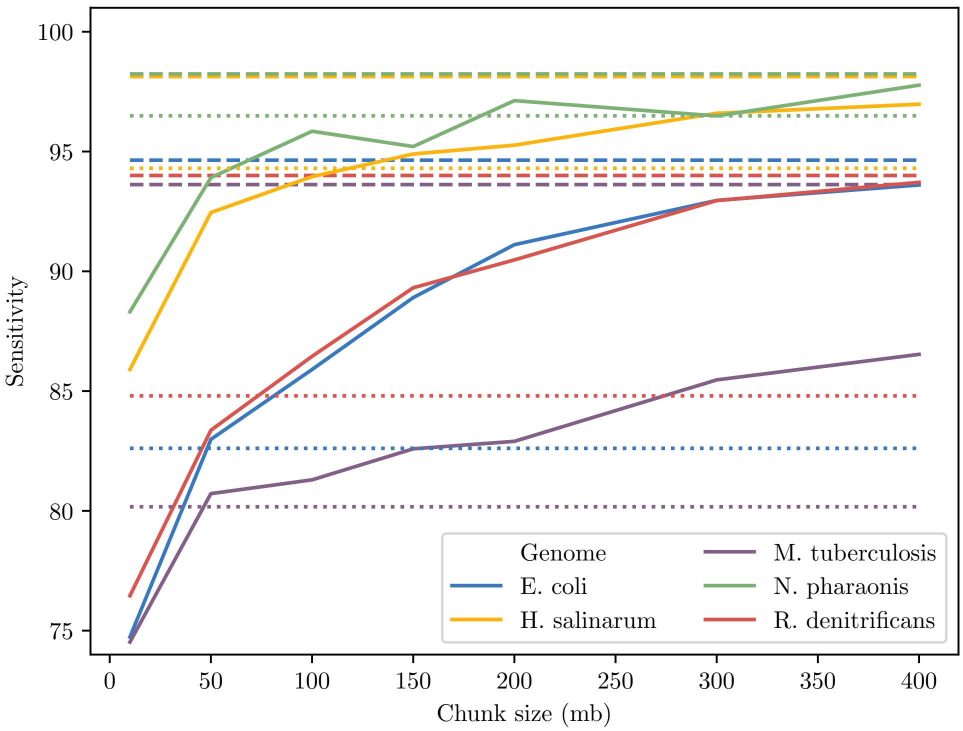


*Fig S9: Gene start prediction sensitivity of GeneMarkS-2 on short fragments of five genomes are shown by solid lines. The sensitivity of MetaGeneMark and StartLink, are shown by the dotted and dashed lines, respectively, for each genome.*

As shown in Fig. S9, the performance of GeneMarkS-2 starts to degrade rapidly when the input fragment size goes below the 200K range. And at 10K, we see a drop all the way down to 75% on *M. tuberculosis*, *R. denitrificans*, and *E. coli*, 96% on *H. salinarum* and 88% on *N. pharaonis*. This is in contrast to StartLink, which doesn't depend on the genome size at all, and maintains an average accuracy of 95% on all genomes, with the lowest being for *M. tuberculosis* at 93.62%.

This experiment shows the advantage of similarity-based approaches in ability to make predictions of gene starts in short sequence fragments for genes whose orthologs are present in the database.

### Note 6: StartLink technical details and optimizations

In this section, we discuss some under-the-hood details of the selection process of target sequences in StartLink, focusing on items that can affect the overall performance and runtime. Before we begin, we state the ultimate goal of this selection process by describing the desired properties of the final selected set of sequences. We then describe the selection process and discuss optimizations and shortcuts made to reduce the overall runtime.

In StartLink, target sequence selection is a multi-step process. We start with a search of protein sequence targets in a database and then filter out unwanted sequences. Given that our initial databases are large, we use DIAMOND BLASTp, which provides a faster version of the BLASTp search algorithm.

This search typically returns a thousands of hits per query, and the goal is to select $N$ (in our case, $N = 50$) target sequences (per query) that will be used by StartLink.

Ultimately, we want to choose a set of sequences that, when aligned, would allow us to find the start of the query gene easily. For our purposes, we want this set $S$ (a query and (maximum) $N$ targets) to satisfy the following pairwise constraints:

$$d_k(s_1, s_2) \in [0.1, 0.5] \qquad \forall s_1, s_2 \in S, s_1 \neq s_2 \tag{13}$$

This means that no pair of sequences in this set should have a Kimura distance smaller than 0.1 or larger than 0.5.

The most straightforward approach to accomplishing this is to compute all pairwise Kimura distances between query and target sequences, and then randomly select $N$ that exist in the desired range. However, the Kimura distance calculation requires a global alignment of two sequences, which in this case would be very costly (given the large initial number of sequences).

#### Avoiding global alignment

Therefore, the first order of approximation is to use the already-existing local alignments between a query and its targets (generated by the BLASTp search), and immediately compute approximate Kimura distance values. In our experiments, we found no tangible difference in StartLink's performance on the set of genes with verified starts based on whether global or local alignment was used in the Kimura distance computation.

Note, however, that these local alignments are only provided between a query and a target, and not between pairs of target sequences. Furthermore, there can still be tens of thousands of targets per query, and going through them all would result in many unnecessary computations.

#### Avoiding all-pairs target alignments

We can avoid (at this stage) directly comparing target sequences to each other by observing how they compare to the query. This step exploits the fact that two *very similar* target sequences should have similar Kimura distances to the query. As such, if the Kimura distance between two targets $t_1$ and $t_2$ and the query is roughly the same (i.e. $d_k(q, t_1) \approx d_k(q, t_2)$), then we remove one of them from the list of candidates under the assumption that there is a higher chance that they are very similar to each other. We deem two Kimura distances as approximately the same if they are equal up to the fourth decimal place.

It's important to note that this heuristic is not optimal, however. In particular, it doesn't hold strictly in the opposite direction; i.e., in general, two sequences with similar Kimura distances to the query may have differences that did not affect the result of computation of the Kimura distance to the query. That said, we've found that this step slightly improves performance because we are less likely at this stage to import multiple target sequences that are very similar to each other, which would end up in biasing the alignment.

#### Avoiding analyzing the large number of target sequences

We first note that hits retrieved from BLASTp are ordered by $e$-value. In practical terms, this means that they are ordered in terms of a similarity score to the query sequence. It also means that they are roughly ordered by the Kimura distance; we say "roughly" because the ordering is not exact, but it does not affect the final performance.

Given this list, we want to find the position of the sequence after which no sequence exists with $d_k \in [a, b]$. If the list was perfectly sorted, this task would boil down to finding the first sequence with $d_k > b$. Given that the ordering is not exact, however, we take some measures to decrease the likelihood that our heuristic removes sequences within the range $[a, b]$.

First, if the total number of target sequences for a query is less than 2,000, we skip this filtering step since the number of computations needed is not large. Second, instead of looking for the first sequence with $d_k > b$, we place an additional buffer that favors keeping in more out-of-range sequences than removing in-range sequences; basically, we update the threshold such that we search for $d_k > b + 0.2$. Third, instead of traversing the list in order and finding the *first* sequence that satisfies $d_k > b + 0.2$, we perform a binary search through the list, which makes it much less likely to stop at a very low positioned sequence in the list.

As such, if the number of sequences is higher than 2,000, we perform a binary search through the list to find an (approximate) "upper bound" above which all sequences (likely) have a distance value larger than the maximum allowed threshold $b$ (+0.2). This operation is quick, requiring $\log(n)$ Kimura distance computations on average (where $n$ is the number of hits). In other words, for a list of 100,000 hits, we require (on average) 16.6 Kimura distance computations, where each computation is a simple run through an *existing* local alignment, at the nucleotide level, counting up the transition and transversion changes.

We effectively now have a list of sequences whose Kimura distance is in the range $[0, b + 0.2]$. We don't perform the same operation to find the lower limit because our desired threshold of 0.1 is close to 0, leaving us a small buffer region. Instead, we shuffle the list and go through it one sequence at a time until we have 50 sequences that satisfy our requirements. In the worst case, we would have to go through all sequences in this range, especially if the target sequences are very similar to the query or each other.

#### Optimality in the face of randomness

If we had gone through every sequence in the list, computed $d_k$, sorted it, then filtered out the sequences outside the range $[a, b]$, we would still have to randomly select only $N$ (from the possibly thousands) of the remaining sequences. This reduction, which comes from the requirement of the subsequent multiple sequence alignment step, means that even if we were to act optimally at the initial stages, the act of random selection reduces the likelihood that our final set is actually optimal. In other words, we can act sub-optimally (as suggested in the above heuristics) at the initial steps without necessarily observing a change in the final outcome. In practice, we did not see a tangible difference in StartLink's final performance by acting optimally or sub-optimally at the initial stages.

### Note 7: Examples of multiple sequence alignments made by StartLink

We show below examples of MSAs where StartLink+ selection of the gene start (#selected) doesn't match PGAP's prediction (#ref). The examples were taken from eight genomes, coming from the four clades considered in the main text.

```
#selected        M-------------------------------------------------
#q-3prime        --------------------------------------------------
#ref             ----------------------M---------------------------
42864;45542;-;A  LstsrhttplqyrfivncilllMisLavcaqsgkntiagrVvdaetsqpv
1;0.3867         Lsidrriysfqyrliscyvlll-isLavyaqsgkerfvgrvvdtetnqpv
2;0.387          Lsidrriysfqyrliscyvlll-isLavyaqsgkerfvgrvvdtetnqpv
3;0.3897         Lsidrriysfqyrliscyvlll-isLavyaqsgkerfvgrvvdtetnqpv
4;0.3925         LnigshiysfqyrLitcyvLll-isLtvfaqsgkerfsgrVidtetnqpv
5;0.3995         LnigshiysfqyrLitcyvLll-isLtvfaqsgkerfsgcVidtetnqpv
6;0.4026         LnidshiysfqyrLiicyvLll-isLtvfaqsgkesfsgrVidtetnqpv
7;0.4038         LnigshiysfqyrLitcyvLll-isLtvfaqsgkerfsgrVidtetnqpv
8;0.405          LnigshiysfqyrLitcyvLll-isLtvfaqsgkerfsgrVidtetnqpv
9;0.4069         LnigshiysfqyrLitcyvLll-isLtvfaqsgkerfsgrVidtetnqpv
10;0.4185        ------MysfqyrLiscyvLll-isLavyaqsgkesfagrvvdtetyqpv
11;0.4201        LnidshiysfqyrLitcyvLll-isLtvfaqsgkerfsgrVidtetyqpv
12;0.4205        LnidshiysfqyrLitcyvLll-isLtvfaqsgkerfsgrVidtetyqpv
13;0.4213        ------MysfqyrLiscyvLll-isLavyaqsgkesfagrvvdtetyqpv
14;0.4226        LnidshiysfqyrLitcyvLll-isLtvfaqsgkerfsgrVidtetyqpv
15;0.423         LnidshiysfqyrLitcyvLll-isLtvfaqsgkerfsgrVidtetyqpv
16;0.4241        LnigshiysfqyrLitcyvLll-isLtvfaqsgkerfsgrVidtetnqpv
17;0.4259        LnidshiysfqyrLitcyvLll-isLtvfaqsgkerfsgrVidtetyqpv
18;0.429         LnidshiysfqyrLitcyvLll-isLtvfaqsgkerfsgrVidtetyqpv
19;0.4292        LnidshiysfqyrLitcyvLll-isLtvfaqsgkerfsgrVidtetyqpv
20;0.4296        LnidshiysfqyrLitcyvLll-isLtvfaqsgkerfsgrVidtetyqpv
```

*Bacteroides reticulotermitis, FCB group*

```
#selected          M-------------------------------------------------
#q-3prime          --------------------------------------------------
#ref               ----------M---------------------------------------
215415;215993;-;A  MrrtvfvaatVlalltagvaaaflagvgpfadtts-adgsdgafptqtta
1;0.1456           Mrrtllvaatvlvlltagisaafltgvgpfadtts-addsdgafptqtta
2;0.146            MrrtllvaatvlvlltagisaafVtgvgpfsddsa--aesdeafptqtta
3;0.164            MrhpllaaatvlallttgvvaafVtgvgpfadttp-agdsdgafptqtta
4;0.1727           MrrtllvaatvlvlltagisaafVtgvgpfsddsa--aesdeafptqtta
5;0.1751           MrrtllvaatvlvlltagvsaafVtgigpfsdda---gdsdepfptqtta
6;0.1773           MrrtllvaatvlvlltagvsaafVtgigpfsdda---gdsdepfptqtta
7;0.1783           MrrtllvaatvlvlltagvsaafVtgigpfsdda---gdsdepfptqtta
8;0.1803           MrrtllvaatvlvlltagisaafVtgvgpfsdds---aesdeafptqtta
9;0.1862           MrrtllvaatvlvlltagvsaafVtgvgpfsddd---aesdepfptktta
10;0.2057          MrrtllvaatvlvlltagvsaafVtgvgpfsddd---aesdepfptktta
11;0.227           MrrtllvaatvlvlltagvsaafVtgigpfsddd---adsdepfptqtta
12;0.2291          MrrtllvaatvlvlltagvsaafVtgigpfsddd---adsdepfptqtta
13;0.2293          MrrtllvaatvlvlltagvsaafVtgigpfsddd---adsdepfptqtta
14;0.2316          MrrtllvaatvlvlltagvsaafVtgigpfsddd---adsdepfptqtta
15;0.3906          MkrsllvasavvllvaagvgvsfVtgigpfadgsdagdqsadpfptqtat
16;0.3976          MkqstlvvlalvalvgsgvgaafVtgvgpfasddd---eldgefptqtat
17;0.4102          MkrstlvvlavlalvgggvatafVtgvgpfasddd---eldgefptqtat
18;0.4142          MkqstlvvlalvalvgggvatafVtgvgpfasddd---eldgefptqtat
19;0.4291          MrrpllavvavllvvgggVtvafatgfgpfaggadssdqptepfptqtpt
20;0.4331          MkrsllvastvvllvaagvgVafVagigpfadgsdagdqstdpfptqtat
```

*Haloferax sp., Archaea*

```
#selected         M-------------------------------------------------
#q-3prime         --------------------------------------------------
#ref              ----------M---------------------------------------
215415;215993;-;A MrrtvfvaatVlalltagvaaaflagvgpfadtts-adgsdgafptqtta
1;0.1456          Mrrtllvaatvlvlltagisaafltgvgpfadtts-addsdgafptqtta
2;0.146           MrrtllvaatvlvlltagisaafVtgvgpfsddsa--aesdeafptqtta
3;0.164           MrhpllaaatvlalllttgvvaafVtgvgpfadttp-agdsdgafptqtta
4;0.1727          MrrtllvaatvlvlltagisaafVtgvgpfsddsa--aesdeafptqtta
5;0.1751          MrrtllvaatvlvlltagvsaafVtgigpfsdda---gdsdepfptqtta
6;0.1773          MrrtllvaatvlvlltagvsaafVtgigpfsdda---gdsdepfptqtta
7;0.1783          MrrtllvaatvlvlltagvsaafVtgigpfsdda---gdsdepfptqtta
8;0.1803          MrrtllvaatvlvlltagisaafVtgvgpfsdds---aesdeafptqtta
9;0.1862          MrrtllvaatvlvlltagvsaafVtgvgpfsddd---aesdepfptktta
10;0.2057         MrrtllvaatvlvlltagvsaafVtgvgpfsddd---aesdepfptktta
11;0.227          MrrtllvaatvlvlltagvsaafVtgigpfsddd---adsdepfptqtta
12;0.2291         MrrtllvaatvlvlltagvsaafVtgigpfsddd---adsdepfptqtta
13;0.2293         MrrtllvaatvlvlltagvsaafVtgigpfsddd---adsdepfptqtta
14;0.2316         MrrtllvaatvlvlltagvsaafVtgigpfsddd---adsdepfptqtta
15;0.3906         MkrsllvasavvllvaagvgvsfVtgigpfadgsdagdqsadpfptqtat
16;0.3976         MkqstlvvlalvalvgsgvgaafVtgvgpfasddd---eldgefptqtat
17;0.4102         MkrstlvvlavlalvgggvatafVtgvgpfasddd---eldgefptqtat
18;0.4142         MkqstlvvlalvalvgggvatafVtgvgpfasddd---eldgefptqtat
19;0.4291         MrrpllavvavllvvgggVtvafatgfgpfaggadssdqptepfptqtpt
20;0.4331         MkrsllvastvvllvaagvgVafVagigpfadgsdagdqstdpfptqtat
```

```
#selected         --------M-----------------------------------------
#q-3prime         --------------------------------------------------
#ref              -----------------------M--------------------------
41253;41750;-;A   --------VklnkiilstlafsvMtsfssfaaydgtptqgeiqlkgelvn
1;0.3436          --------MktnklaiiaalaagMtsMtafaaytgspttgeiqfqgelvn
2;0.3439          --------MqtnklviiaalaagMtsMtafaaytgspttgeiqfqgelvn
3;0.3755          --------MkMnkvaMavaftaaMssMsvla----dttngqiefqgeLvn
4;0.3887          ywykqdivMklnkvaMavaftaalgsMsvla----dttngqiefqgelVn
5;0.3915          -LvktgyfMklnkvaMavaLtaalgsMsafa----dttngqiefqgelVn
6;0.4049          --------MkLnkvaMavaftaalgstsvla----antngtiefqgelvn
7;0.4157          ywykqeivMkMnkvaMavafsaalgsMsvla----dttngViefqgelvn
8;0.4235          nLykqdivMnMnkvaMavafsaalgsMsvlaa---dttngMiefqgelvn
9;0.4302          --------MkLnkvalavaftaavssMsvla----dttngqiqfqgeLVn
10;0.4346         lwykqdivMkMnkvaltvaftaalssasvfa----attngqiefqgelVn
11;0.4358         ywykqdivMkMnkvaMavaftaalgsMsVla----dttngViefqgelVn
12;0.4392         --------MkLnkvalavaftaavssMsvla----dttngqiqfqgeLVn
13;0.4457         --------MklnkvaMaValtaalgsMsala----dttngqielqgelVn
14;0.4622         iwykqdivMkMnkvaMavaftaalssMsvlaa---dtttgkiefqgelVn
```

*Tatumella saanichensis, Enterobacterales*

```
#selected          ----------------------M---------------------------
#q-3prime          --------------------------------------------------
#ref               ----------------------------------------M---------
234115;235320;-;A  ----------------------VtapnglrtiptgdrawlMaalgdalsg
1;0.1229           ----------------------VtapnglrtiptddrawlMaalgdaLsg
2;0.1328           ----------------------VtapnglrtiptddrawlMaalgdalsg
3;0.2258           d----rshgarpVhggaeypgrVtsphrlraiptedrawLtaaladalhg
4;0.2606           --------VfgpdpatagilegVtskapiraiptgdrawllaaladaldg
5;0.2782           ppcaaparrrcaqqgcaeypgrVtaphrlraiptedrawltagladaLrg
6;0.2841           ----------------------Vtaphvlrtipvedrawliaalddalag
7;0.2896           ------------tpsereyaewVtaphglraiptedrawlMagladalag
8;0.328            ----------------------Vtsphglriipvsdrawlmaalgdalag
9;0.3689           ---------prppltearyprwVtaplvlraipvedrawliaalddalag
```

*Leucobacter chironomi, Actinobacteria*

```
#selected          M-------------------------------------------------
#q-3prime          --------------------------------------------------
#ref               ---------------------M----------------------------
4859339;4862848;+;A MkkrwksnfplgrklqkifrcMkltfllltcfvvqtfavlnaqtvtikkq
1;0.1135           MkkrwksdfplerkLqkifrcMkltfllltcfvvqtfaslnaqtvtikkq
2;0.1176           MkkrwksdfplerkLqkifrcMkltfllltcfvvqtfaslnaqtvtikkq
3;0.2777           MekrwknvfpsggntrkMircMkltLllltcfVlqtfaganaqivtikkq
4;0.3134           MkkrwksdfplrgkLqkiircMkltfllltcfVvqtfaslhaqtvtikkq
5;0.3306           MkkrwksifpsggntrkMircMkltLllltcfVlqtfaganaqtvtikkq
6;0.3453           MkkrwksvfpsggnMrkMircMkLtlllltcfViqtfaganaqtvtikkq
```

*Butyricimonas sp., FCB group*

```
#selected          -M------------------------------------------------
#q-3prime          -*------------------------------------------------
#ref               --------------------------------------------------
20902;21885;-;B    LMesrseqfkndifeisaeagdarsgvlqigdtklstpnllpvvnfyagg
1;0.124            VMesrseqfkndifeisaeagdarsgslqigdtklstpnllpvvnfyagg
2;0.1547           VMesrseqfkndifeisaeagdarsgvLrigdtelstpnllpvvnfyagg
3;0.1756           MassgsqrlddgifeiideagdarsgVisindteletpnLlpvVnfyagg
4;0.1777           MassgsqrlddgifeivdeagdarsgVisindteletpnLlpVVnfyagg
5;0.1927           MapsgsqrlddglfeiiseagdarsgVlriddteLetpnllpVVnfyagg
6;0.1964           MthsgsprldddLfeitseagdarsgVirfddteletpnLlpVVnfyagg
7;0.4616           -MsfdipetkvasfevdstvgdaragtlrikdteletpnllpVVnfyagg
8;0.4712           -MpfssprtdiatfdiestagdaragtlriedtslqtpnllpvVnlyagg

#selected          --------------------------------------------------
#q-3prime          --------------------------------------------------
#ref               -------------------------------------M------------
20902;21885;-;B    tdaslygggihrtikefMngdevvnggdysryfdgVMtsvasltdygisr
1;0.124            tdaslygggihrtikefMngdevvnggdysryfdgVMtsvasltdygisr
2;0.1547           tdaslygggihrtikefMngdevvnggdysryfdgvMtsvasltdygisr
3;0.1756           tdsslygggihrtikefMngdevinggdysryfdgVMtsvasltdygisr
4;0.1777           tdsslygggihrtikefMngdevinggdysryfdgVMtsvasltdygisr
5;0.1927           tdsslygggihrtikefingddvvnggdysryfdgVMtsvssltdygisr
6;0.1964           tdsslygggihrtikefingdevvnggdysryfdgVMtsvasltdygisr
7;0.4616           larslygggihrtMkefMtgddviggddyseyfdgvMtsvgsltdynisr
8;0.4712           MdrslygggihrtMkefMtghdvigggdyseffdgiMtsvgsltdynisr
```

*Halorubrum kocurii, Archaea*

```
#selected          ---------M----------------------------------------
#q-3prime          --------------------------------------------------
#ref               ---------------------------------M----------------
386604;389732;+;A  ---------MpnklptafsritrssgransiltVtlplLLvliatlfwsa
1;0.1949           ---------Mpnksptafsritrssgrtnsiltvtlpllfvlittlfwsa
2;0.197            ---------Mpnksptafsritrssgrtnsiltvtlpllfvlittlfwsa
3;0.4132           ---------Manssptafsritrnsgrtnsiltvalpllfiliatlvwsa
4;0.4284           LpiidensaMstpsptafsritrtsgrtnsiltiafpLlfilittLiwsa
```

*Providencia alcalifaciens, Enterobacterales*

```
#selected          -------------------M------------------------------
#q-3prime          --------------------------------------------------
#ref               M-------------------------------------------------
3718301;3719053;+;C MprpgvpplthrtrvyperVtssasagfrtrpehrwpVlialgvgtgLya
1;0.1864           acspraatrrsrtqvypepVstheregfrtrpehrwpvlialavgtglya
2;0.1885           -----MssirdraeeyperVnthdragfrtrpehrwpVlialaVgtglya
3;0.2015           -------------------VspqdpatfrtrpehrwpVlialavgtglya
4;0.2058           -------------------VtssdradfrtrpehrwpVlislglgtilya
5;0.208            -------------------VstheregfrtrpehrwpvLialavgtglya
6;0.2081           pfrahgekpthrpgvyperVnqrqqgafrtrpehrwpVlialaVgaglya
7;0.2103           -------------------VspqdpaafrtrpehrwpVvialavgaglya
8;0.2147           -------------------Vstheregfrtrpehrwpvlialavgtglya
9;0.2148           -------------------VsrqdpsafrtrpehrwpVlialaVgaglya
10;0.2169          -------------------VstqdaagfrtrpehrwpVlialavgaglya
11;0.2795          -------------------VtskesaefrtrpehrwpVlifLalgtVLya
12;0.4002          ----------------------MtaadfrarpehrwpvvvallVaVVlyl
13;0.4759          ----------------------VattefrsraehrwpvVvaiaVaialyl
14;0.4783          -----------------------Maqqrrsaaeyrwppvlaivvalalya
15;0.4809          -VtrqapMlrhtlglM------eqhpsrqVpaehrwpaiigVvaalalyt
16;0.4831          ---------------V------pihpsrrtvaeprwpaavglivavalyg
17;0.4849          ---------------V------pihpsrrtvaeprwpaavglvvavvlya
18;0.4865          -------------------------------------------lhnglva
19;0.4874          --------------VV------tqhpsrtirpehrvpaliavvaalllys
20;0.4911          ---------------V------pihpsrrtvaeprwpaavglvvavvlya
```

*Microbacterium testaceum, Actinobacteria*
